# Transient let-7/miR-98 reprograms primed pluripotency to induce naive human PSCs

**DOI:** 10.1101/2025.10.14.682465

**Authors:** Yoshihiko Fujita, Moe Hirosawa, Karin Hayashi, Hiroki Ono, Belinda Kaswandy, Mai Ueda, Maya Lopez, Takuya Yamamoto, Yasuhiro Takashima, Hirohide Saito

## Abstract

Naive human pluripotent stem cells (hPSCs) are valuable for modeling early development and enabling regenerative applications, yet resetting from the primed state remains inefficient and mechanistically unclear. We identify an unexpected role for miR-98-5p, a let-7 family microRNA (miRNA) typically linked to differentiation, in promoting naive induction. Using a synthetic mRNA screen containing diverse miRNA-responsive elements, we uncovered miR-98-5p as a potent enhancer of resetting. Single-cell RNA sequencing and targeted perturbations show that miR-98-5p directly represses NR6A1 while transiently activating BMP2/4 signaling, triggering network transitions. These changes destabilize the primed gene-regulatory network and enable acquisition of a naive state. Leveraging this mechanism, we establish a feeder-free, miRNA-based, editing-free protocol that boosts resetting efficiency from <10% to ∼80%. To our knowledge, this is the first demonstration that a let-7 miRNA can drive naive induction by coupling direct target repression with activation of pro-naive signaling, suggesting a generalizable reprogramming principle of transient network destabilization.

## Introduction

Embryonic stem cells (ESCs) are derived from fertilized eggs during the blastocyst stage, and numerous studies have collectively demonstrated various species-specific differences in ESC properties between mice and humans. Mouse ES cells (mESCs) develop a dome-like shape, while human ES cells (hESCs) form flat colonies.^1^ hESCs possess comparable morphology and gene expression profiles to epiblast stem cells (EpiSCs) derived from mouse post-implantation embryos. Thus, conventional hESCs and mESCs are described as primed and naïve pluripotent stem cells, respectively.^2^ Naïve human pluripotent stem cells (naïve hPSCs) exhibit morphological and molecular similarities with mESCs. Recent studies have shown that naïve hPSCs have the potential to differentiate into early developmental cell types, including trophoblasts.^3^ Additionally, naïve hPSCs can reset the epigenomic state, and, as a result, re-primed human induced pluripotent stem cells (hiPSCs) generated from naïve hPSCs may lose their differentiation bias, enabling them to differentiate into various cell types more efficiently.^4^ Given these advantages, naïve hPSCs have become a critical model for studying early human development and differentiation, while also representing a valuable resource for regenerative medicine and tissue engineering.^5,6^ Thus, investigating the reprogramming pathway and developing robust and efficient production methodologies of naïve hPSCs are imperative to harness their potential.

The transformation of primed hESCs/hiPSCs towards naïve hPSCs, termed “resetting,” can be facilitated by the forced expression of key transcription factors, such as Nanog and Klf2.^7^ In addition, a method for generating naïve hPSCs using small molecules has been reported.^8–10^ In these methods with chemical compounds, primed hESCs are treated with HDAC inhibitors (e.g., valproic acid; VPA), presumably causing widespread changes in gene expression through epigenomic change. While multiple signaling pathways, such as WNT and MEK, are known to be involved in naïve pluripotency,^11^ the mechanistic details of how they are involved in resetting primed hESCs to naïve hPSCs are not fully understood. The expression of Oct4, Sox2, Nanog, and other genes in primed cells forms a stable state by interacting with each other in a complex manner.^12^ Destabilization of the gene networks and acquisition of cellular plasticity, together driving the switch toward the naïve network, are assumed to be essential during the conversion from the primed to naïve state. However, conventional approaches to reprogram hESCs or hiPSCs into the naïve state remain inefficient, and the underlying molecular mechanisms are poorly understood.

MicroRNAs (miRNAs) are ∼22-nucleotide non-coding RNAs that simultaneously regulate multiple gene expression events.^13,14^ In humans, approximately 2,600 distinct miRNAs are registered in miRbase,^15^ and many exhibit dynamic expression changes during development, contributing substantially to cellular functions. For example, the let-7 family—one of the best-characterized miRNA families—has been highly conserved through evolution and is known to promote differentiation by repressing stemness-associated genes.^16–18^ The miR-302/367 cluster has been reported to enhance reprogramming efficiency in generating hiPSCs from human fibroblasts.^19^ Moreover, several studies have shown that miRNAs highly expressed in specific target cells can improve the differentiation efficiency of mouse or human iPSCs/ESCs into diverse lineages, including neurons, endothelial cells, and hepatocytes.^20–22^ Based on these findings, we hypothesized that additional, yet unidentified, miRNAs might facilitate the transition from primed to naïve states.

In this study, we unexpectedly discovered that human miR-98-5p, a member of the let-7 miRNA family—widely recognized for promoting differentiation and suppressing self-renewal in pluripotent stem cells^17,23^—instead facilitates the resetting of hESCs/hiPSCs into naïve hPSCs. Building on this counterintuitive observation, we investigated to elucidate how miR-98-5p enhances resetting efficiency. Our analyses revealed that Nuclear Receptor subfamily 6 group A member 1 (NR6A1) and Bone Morphogenetic Protein 2 (BMP2) are negatively and positively regulated by miR-98-5p, respectively, and are critical for efficient naïve-state induction. Importantly, we identified transient, optimally timed BMP2/4 signaling during the early resetting phase as essential for naïve hPSC generation. Leveraging this insight, we substantially improved the efficiency of naïve hPSC generation under feeder-free culture conditions. This discovery establishes a powerful strategy for regulating human cell reprogramming and offers new perspectives for regenerative medicine and synthetic biology.

## Results

### MicroRNA-sensing mRNA library identifies miR-98-5p as a driver of naïve hPSC conversion

The potential of miRNAs to direct cellular differentiation and reprogramming has been underscored by several studies demonstrating that cell type–specific miRNAs can mediate cell fate conversion.^19,21,24^ Building on these findings, we investigated miRNA activity profiles between naïve hPSCs and primed hESCs by developing a miRNA-sensing reporter mRNA library (Figure 1A). We first generated a library of 720 distinct miRNA-responsive mRNAs by inserting complementary sequences to target miRNAs—defined as confident miRNAs in miRBase^15^—into the 5′ untranslated regions (5′ UTRs).^25–27^ Endogenous miRNAs bind to their complementary sequences and repress reporter expression via Argonaute 2–mediated regulation,^25^ enabling quantification of each miRNA’s activity in cells. For this screen, we used the hESC line, H9-NK2,^6^ which carries an EGFP cassette downstream of the EOS (Early transposon promoter with Oct-4 [Pou5f1] and Sox2 enhancers) element, allowing EGFP expression upon conversion to the naïve state.^28^

**Figure 1.**
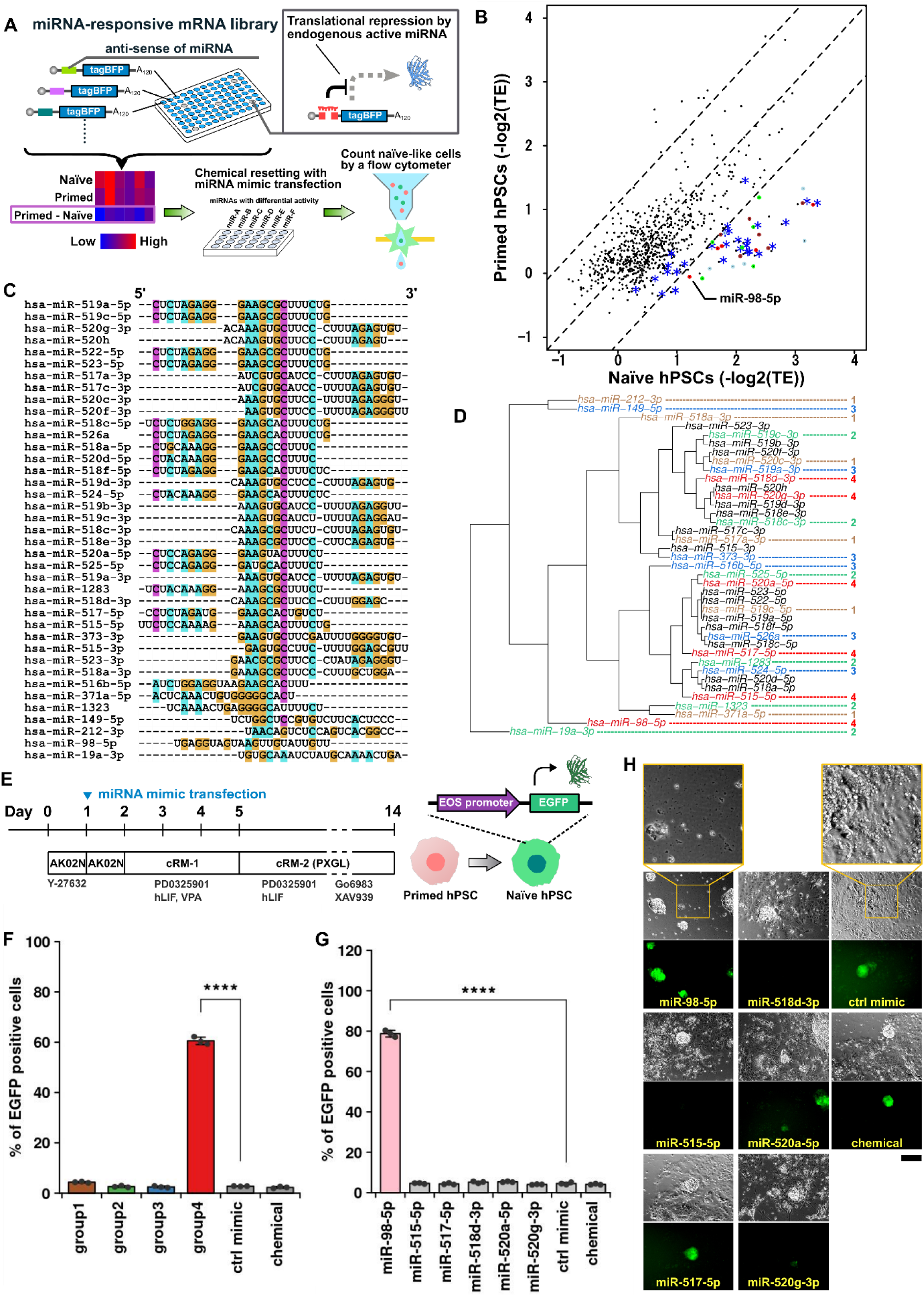
Identification of miR-98-5p as a miRNA driver of naïve hPSC conversion. A: Schematic illustration of the miRNA activity screening. Approximately 720 mRNAs containing different miRNA-complementary sequences were introduced into the 5′-UTR of tagBFP, generating miRNA-responsive OFF switches. Expression of tagBFP from these constructs was suppressed by the corresponding endogenous miRNAs. The degree of suppression of tagBFP expression was defined as miRNA activity in naïve hPSCs and primed ESCs. Candidate miRNAs were selected based on differential activity between the two states, then transfected during the resetting process from primed hPSCs (hESCs or hiPSCs) to naïve hPSCs, and assayed for their ability to enhance resetting efficiency. B: miRNA activity profile of naïve and primed hPSCs. The rate of decrease in fluorescence of the miRNA-responsive OFF switch relative to the normal tagBFP mRNA is defined as activity and plotted. Blue dots indicate miRNAs previously reported as highly expressed in naïve hPSCs. Brown, green, light blue, and red dots indicate miRNAs corresponding to groups 1 to 4, respectively, described in Figure 1D. ‘*’ indicates highly expressing miRNAs in naïve hPSCs. Dashed lines show log2 fold change is -1.1, 0, and 1.1, respectively. C: Alignment of differentially active miRNAs in naïve hPSCs. ClustalW was used to align miRNAs with 2^1.1^-fold higher activity in naïve hPSCs than primed hPSCs. D: Dendrogram of aligned miRNAs. Alignment results were displayed in a dendrogram, with miRNAs grouped according to their sequence similarity, as indicated by the numbers (1-4) to the right of each miRNA. miRNAs with black letters were excluded from further analysis. E: Typical resetting procedure with miRNA under chemical resetting. miRNA mimics were transfected after replacing the medium with AK02N on day 1. Cells were then cultured in cRM-1 medium containing VPA for 3 days (from day 2), followed by culture in cRM-2, a naïve maintenance medium. H9-NK2 and H9EOS cells harbor EGFP under the control of the EOS promoter in their genomes. EGFP-positive cells observed on day 14 were counted as naïve-like hPSCs. F: Enhancement of resetting efficiency in H9-NK2 cells by groups of different miRNA cocktails. Based on the dendrogram in D, miRNAs were divided into four groups and transfected into cells. The resetting efficiency (percentage of EGFP-positive cells) was measured on day 14. Technical replicate (n=3). \*\*\*\**P* = 2.07×10^-4^. G: Enhancement of resetting efficiency of H9-NK2 cells by each miRNA in group 4. Group 4 miRNAs were transfected separately, and the resetting efficiency was measured. Technical replicate (n=3). \*\*\*\**P* = 3.69×10^-5^. H: Microscopic images of H9-NK2 cells treated with each miRNA in group 4 on day 14. Representative loss of non-naïve cells following transfection with miR-98-5p is shown enlarged at the top, whereas control mimic–treated cells retained non-naïve cells. Scale bar, 200 μm.

These 720 distinct miRNA-responsive mRNAs were introduced into either primed hESCs or naïve hPSCs generated via a chemically induced, feeder-based protocol.^8^ Reporter gene expression, quantified as *−log₂(translational efficiency)*, was used to estimate miRNA activity in each cell type (Figure 1B). We validated the reporter library’s ability to capture miRNA activity profiles by examining several miRNAs—miR-371 and members of the miR-518 and miR-520 families—known to be highly expressed in naïve hPSCs (Figure 1B, dots marked by ‘*’).^24,25^ We reasoned that profiling miRNA activity, rather than relying solely on miRNA expression levels, would provide a more direct approach to identifying novel miRNAs capable of driving naïve hPSC reprogramming, as miRNA expression does not always reflect functional activity.^26,27^ To our knowledge, miRNA activity had never before been systematically assessed in the context of naïve hPSCs.

Among the 720 miRNAs tested, we selected 39 candidates from those that reduced reporter expression (reflecting higher miRNA activity) in naïve hPSCs relative to primed hESCs. To refine this set, we performed sequence alignment and clustering analysis, discarding miRNAs with identical or highly similar sequences (Figure 1C). The remaining 24 sequences were grouped into four distinct clusters based on low sequence similarity (groups 1–4) (Figure 1D). To assess their effects during resetting, we prepared cocktails of synthetic miRNA mimics corresponding to each group and transfected them into hESCs (H9-NK2 line) on day 1. Resetting from the primed to the naïve state was initiated by culturing cells in cRM-1 medium including valproic acid (VPA), a histone deacetylase inhibitor that destabilizes epigenetic marks in the chemical resetting protocol.^8,9^ Cells were subsequently switched to cRM-2 (PXGL medium), a maintenance medium for naïve hPSCs containing Go6983 (PKC inhibitor) and XAV939 (WNT inhibitor). The number of EGFP-positive cells was measured by flow cytometry on day 14 (Figure 1E).

Notably, group 4 miRNAs significantly increased the number of EGFP-positive cells compared to the control mimic, whereas the other three groups had no effect (Figure 1F). Detailed analysis using individual transfections of group 4 mimics revealed that miR-98-5p increased the number of EGFP-positive cells by ∼80%, compared to <10% for other candidates (Figure 1G, H). We also observed that miR-98-5p may preferentially eliminate cells failing to acquire a naïve-like state during the early resetting phase (Figure 1H, magnified inset).

The H9-NK2 hESC line was genetically engineered to carry a Dox-inducible *NANOG* or *KLF2* transgene cassette, although these genes were not induced in this study^7^ (Figure 1E–G). Nevertheless, we considered the possibility that leaky *NANOG* or *KLF2* expression (in the absence of Dox) might influence resetting efficiency. To exclude this potential effect, we performed resetting experiments with another hESC line, H9EOS, which does not contain the *NANOG* or *KLF2* cassette. The results confirmed that the miR-98-5p mimic significantly increased the number of EGFP-positive cells, whereas other miRNA mimics—including those from the miR-371, miR-518, and miR-520 families, which are highly expressed in naïve hPSCs (blue dots in Figure 1B)—did not (Figure 2A). These findings suggest that, unlike the generation of primed hiPSCs driven by the miR-302 family,^19^ which is highly expressed in primed hiPSCs/ESCs, the miR-371/518/520 families do not induce naïve hPSCs under the tested conditions. This further implies that forced expression of miRNAs enriched in target cells does not necessarily lead to cell fate conversion.

**Figure 2.**
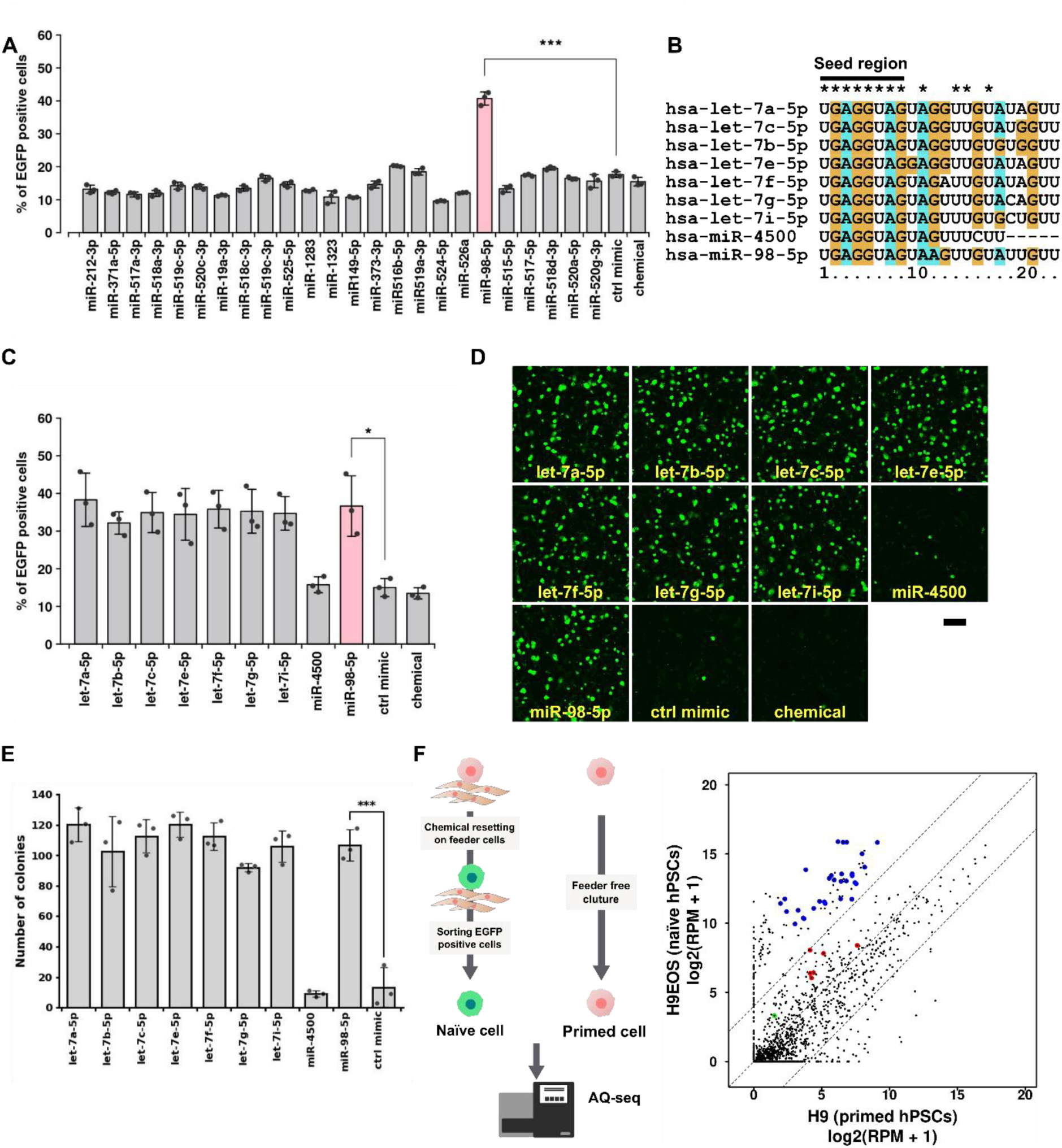
miRNA-driven resetting of H9EOS cells from primed to naïve state. A: Enhancement of resetting efficiency by transfecting each candidate miRNA on day 1. Biological replicate (n=3). \*\*\**P* = 6.30×10^-4^. B: Alignment of miRNAs whose seed sequence (8 mer from 5’end) matches miR-98-5p. ‘*’ indicates the same nucleotide as miR-98-5p. C: Resetting efficiency of let-7 family miRNAs whose seed sequence matches miR-98-5p. Biological replicate (n=3). \**P* = 3.42×10^-2^. D: Fluorescent microscopy images of the H9EOS cells transfected with each miRNA on day 14. Scale bar, 1 mm. E: Quantification of naïve-like colony numbers on day 14 for each miRNA. Colony numbers were obtained by automated image analysis of the fluorescence images in **D**. Biological replicate (n=3). \*\*\**P* = 8.10×10^-4^. F: A miRNA expression profile of naïve H9EOS cells compared to primed H9 cells. H9EOS cells were used and sorted based on EGFP expression by a cell sorter to purify naïve hPSCs. The blue, red, and green dots indicate miRNAs highly expressed in naive cells, miRNAs of the let-7 family, and miR-98-5p, respectively. Biological replicate (n=2). Dotted lines indicate log2 fold changes of ±4.

Given the crucial role of the miRNA seed sequence in mRNA recognition, we searched miRBase^15^ for mature miRNAs sharing the same seed sequence as miR-98-5p. Most of these miRNAs belonged to the let-7 family (Figure 2B). Interestingly, transfection of H9EOS cells with mimics of these let-7 family miRNAs—except for miR-4500—resulted in resetting efficiencies comparable to that of miR-98-5p (Figures 2C, D). To verify that this effect reflected an actual increase in the number of naïve-like colonies, rather than merely a higher proportion, we quantified colony numbers by image analysis. This revealed a clear rise in the absolute number of naïve-like colonies (Figure 2E, S1A), indicating that the let-7 miRNA family broadly facilitates the resetting of hESCs into naïve hPSC-like cells.

Next, we analyzed the expression levels of miRNAs, including miR-98-5p and other let-7 family members in both naïve and primed H9EOS cells using AQ-Seq,^29^ which quantitatively detects miRNA expression. We sorted EGFP-positive naïve hPSC-like cells after resetting at day 14 and compared their miRNA profiles with those of primed hESCs. We observed clear upregulation of naïve marker miRNAs such as miR-371 and miR-518 (blue dots in Figure 2F), whereas miR-98-5p and other let-7 family members (red dots) remained largely unchanged compared to naïve marker miRNAs. These results suggest that the transient gene regulation mediated by let-7 family miRNA mimics increases the number of EGFP-positive naïve hPSC-like cells without altering steady-state expression levels.

Furthermore, to determine the stage during which transient miR-98-5p activation enhances resetting, we investigated the effects of transfection timing. Improved efficiency was observed when miR-98-5p was introduced just before cRM-1 treatment (day 1), whereas repeated transfection on days 1 and 2 did not further increase the number of EGFP-positive cells in either H9EOS or H9-NK2 line. This indicates that a brief, early pulse of miR-98-5p activity is sufficient to promote naïve-state induction.

### Single-cell RNA-seq analysis identifies specific gene sets regulated by miR-98-5p

To examine how miR-98-5p promotes the conversion from primed to naïve states, we performed single-cell RNA sequencing (scRNA-seq) on cells transfected with either an miR-98-5p mimic or a negative control mimic lacking interaction with any mRNAs. Repeated introduction of miR-98-5p mimic did not further enhance resetting efficiency compared with a single introduction on day 1, suggesting that the critical effects of miR-98-5p occur shortly after initial introduction. Given that miRNA activity generally becomes evident within 24-48 hours, we therefore analyzed cells at day 3 (48 hours post-transfection) to capture these early events. We collected cells on days 3, 8, and 14 after transfection with miR-98-5p or control mimics for scRNA-seq, as EGFP-positive cells first appeared on day 8 during resetting (Figure 3A, B). By day 14, a distinct cell population emerged in the miR-98-5p-transfected group, characterized by increased expression of naïve markers (e.g., *DNMT3L*, *DPPA3*, *DPPA5*, and *KLF5*) and decreased expression of primed hESC markers (e.g., *CD24*, *ZIC2*, and *SFRP2*) (Figure 3C, D; Figure S2A, B).^28,30^ These findings indicate that EGFP-positive cells had acquired a naïve hPSC-like state and that the resetting process was actively underway.

**Figure 3.**
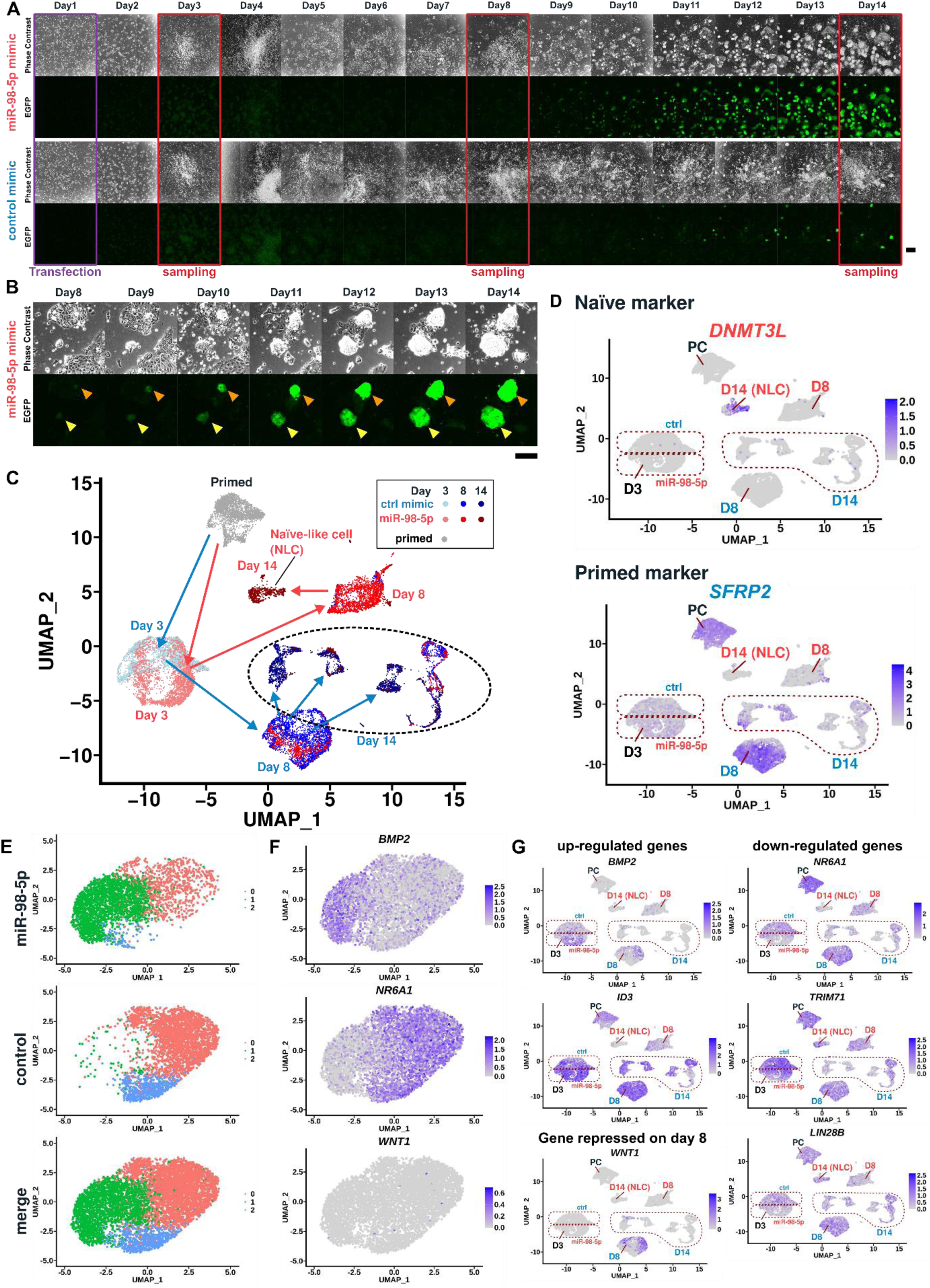
scRNA-seq analysis of miR-98-5p–induced changes during resetting. A: Time-lapse images of the resetting process with miR-98-5p- or control (ctrl) mimic. A miRNA mimic was transfected on day 1 after medium replacement. Cells indicated by red rectangles were harvested and analyzed for scRNA-seq. Scale bar, 200 μm. B: Magnified time-lapse images at the same position after day 8. Orange and yellow arrowheads indicate the same position, respectively. Scale bar, 200 μm. C: UMAP analysis of gene expression in primed cells and cells at days 3, 8, and 14 during resetting. Labels indicate sample identities predominantly included in each cluster. Red and blue arrows connect miRNA-98-5p- or ctrl mimic-transfected cell populations, respectively, across time points. D: Expression levels of naïve and primed marker, *DNMT3L* and *SFRP2*. Additional genes are shown in Figure S2. E: Clustering analysis of cells in Cluster 0 in Figure S3B (resolution = 0.2). Cells transfected with ctrl or miR-98-5p mimics are displayed separately or merged. Cells were separated into three groups. Dot colors indicate cell clusters. Ctrl mimic-transfected cells were predominantly observed in pink and light-blue clusters, whereas miR-98-5p-transfected cells were mainly in the green cluster. F: Representative expression pattern of differentially expressed genes (*BMP2* and *NR6A1*) and a suppressed gene (*WNT1*) in Cluster 0, derived from miR-98-5p–treated cells (Figure S3B). *BMP2* and *NR6A1* were expressed exclusively in the green cluster or primarily in the red cluster, respectively. G: Global expression levels of differentially expressed marker genes on day 3. NLC and PC indicate naïve-like cell and primed cell, respectively.

Interestingly, we observed a modest transcriptomic shift between cells treated with miR-98-5p and control mimics on day 3 (Figure 3C, light blue and pink). To examine whether this discrepancy arose earlier, we additionally analyzed cells collected on day 2 but found no significant differences between the main populations transfected with miR-98-5p or control mimics (Figure S3A). We then performed clustering across all samples and identified 10 distinct clusters (Figure S3B). The velocity-informed latent time increased monotonically with sampling days (ctrl day 0 < day 3 < day 8 < day 14), consistent with the actual sampling time points (Figure S3C). Although velocity analysis did not reveal clear connections between distant clusters, likely due to their wide separation in the embedded space, it uncovered locally divergent flow patterns within individual clusters (Figure S3D). Specifically, cluster 0, composed primarily of day 3 cells, exhibited two distinct flow directions, suggesting early divergence in cell fate. Similarly, clusters enriched for day 8 cells treated with either ctrl mimic or miR-98-5p displayed multiple flow directions, indicating that a subset of cells may not fully progress toward the naïve-like state. These findings are consistent with the incomplete conversion efficiency observed during naïve induction.

To further analyze day 3 cells, we extracted and re-clustered Cluster 0 because cells on day 3 are primarily located within Cluster 0 (Figure S3B) as identified by UMAP. As a result, Cluster 0 was separated into three clusters (Figure 3E), with clear segregation of miR-98-5p- and control mimic-transfected cells. Upon comparison of these day-3 populations, we observed that miR-98-5p transfection led to the upregulation of several genes, such as *BMP2* and *ID3*, and downregulation of others, including *NR6A1*, *TRIM71*, and *LIN28B* (Figure 3F, G). Additionally, by day 8, specific genes of interest, like *WNT1*, were suppressed in cells transfected with miR-98-5p (Figure 3F, G, Figure S2C, D). Notably, some are well-known genes associated with a wide range of differentiation processes, such as maintaining pluripotency (*LIN28*),^31^ driving neural differentiation (WNT signaling),^32^ and promoting differentiation into trophoblasts (BMP signaling).^33,34^

Next, to examine intermediate cell types and transcriptional changes during the transition from the primed to the naïve state, we compared gene ontology (GO) profiles of the top 50 upregulated genes in each cluster (Figure S3E, F, G, H). Control mimic–transfected cells (Cluster 1, day8 and Cluster7, day 14) predominantly upregulated genes related to neural differentiation, including WNT signaling and midbrain development, suggesting a tendency toward neural lineages rather than naïve reprogramming. In contrast, miR-98-5p–treated cells (Cluster 3, day 8) showed enrichment of muscle-related gene signatures, although cell type assignment remained ambiguous due to high p-values. These results indicate that, under cRM medium conditions (Ndiff-based neural differentiation medium), in the absence of miR-98-5p, most cells lose pluripotency and differentiate toward neural fates, thereby inhibiting resetting to the naïve state.

Gene set enrichment analysis (GSEA) further revealed that, compared with control mimic-treated cluster (Cluster 1), miR-98-5p–treated cells (Cluster 3) activated WNT/β-catenin transcriptional pathways and AP1/ZEB1/DLX target genes, while repressing lineage-stabilizing networks such as GATA1/RFX1 (see Table S2). Notably, although WNT1 expression was reduced in Cluster 3 (shown as “D8” in Figure 3G), enrichment of CTNNB1-dependent and FZD8-CRD–blocked WNT gene sets is consistent with pathway-level activation, potentially mediated by alternative WNT ligands such as WNT6 (Figure S3J). Together, these findings suggest that miR-98-5p promotes naïve reprogramming by redirecting transcriptional programs away from neural differentiation and toward pathways permissive for naïve-state acquisition.

To further characterize each cluster, we performed automatic cell-type assignment using ScType^35^ with marker gene sets from the CellMarker 2.0 database^36^. Clusters 1 and 7 (control mimic–treated cells) were classified as neural cells, while clusters 5, 8, and 4 were annotated as neural-associated cells (Figure S3I), consistent with the GO analysis and supporting the reliability of our approach. Cluster 3 (miR-98-5p–treated cells) was assigned as trophectoderm, consistent with the upregulation of several trophectoderm-related genes (*KRT19, KRT8, KRT18, CD24, and PDLIM1*)^37^ (Figure S3J, K). Previous studies have reported that primed-state cells can transition to amnion.^6,38^ Accordingly, we also examined amnion markers and observed upregulation of related genes (e.g., *GABRP, ISL1, and WNT*6)^39^ (Figure S3J, K). Clusters 0 and 5 were assigned to Sertoli cells and ciliated epithelial cells, respectively; however, only minor differences in marker gene expression compared to other clusters limited the reliability of these annotations (Figure S3K). Overall, our scRNA-seq analysis suggests that the miR-98-5p mimic enhances resetting efficiency, particularly during the early phase, by activating pathways such as BMP signaling while suppressing others, including WNT1, that may influence progression toward the naïve state.

### Transient BMP signal plays a critical role in the resetting process

To identify genes contributing to resetting efficiency, we constructed Dox-inducible *piggyBac* expression vectors for 22 genes upregulated by miR-98-5p treatment. These genes were differentially expressed, both in fold change and in the proportion of expressing cells, between miR-98-5p– and control mimic–transfected cells in Cluster 0 (≈ day 3) (Figures 3E, F, S3). H9EOS cells were transfected with each plasmid in the presence of Dox, and resetting efficiency was assessed by the proportion of EGFP-positive cells at day 14. Transfection with a mixture of all 22 plasmids (all-mix) significantly promoted resetting, whereas individual plasmids alone were largely ineffective (Figures 4A, B, S4).

**Figure 4.**
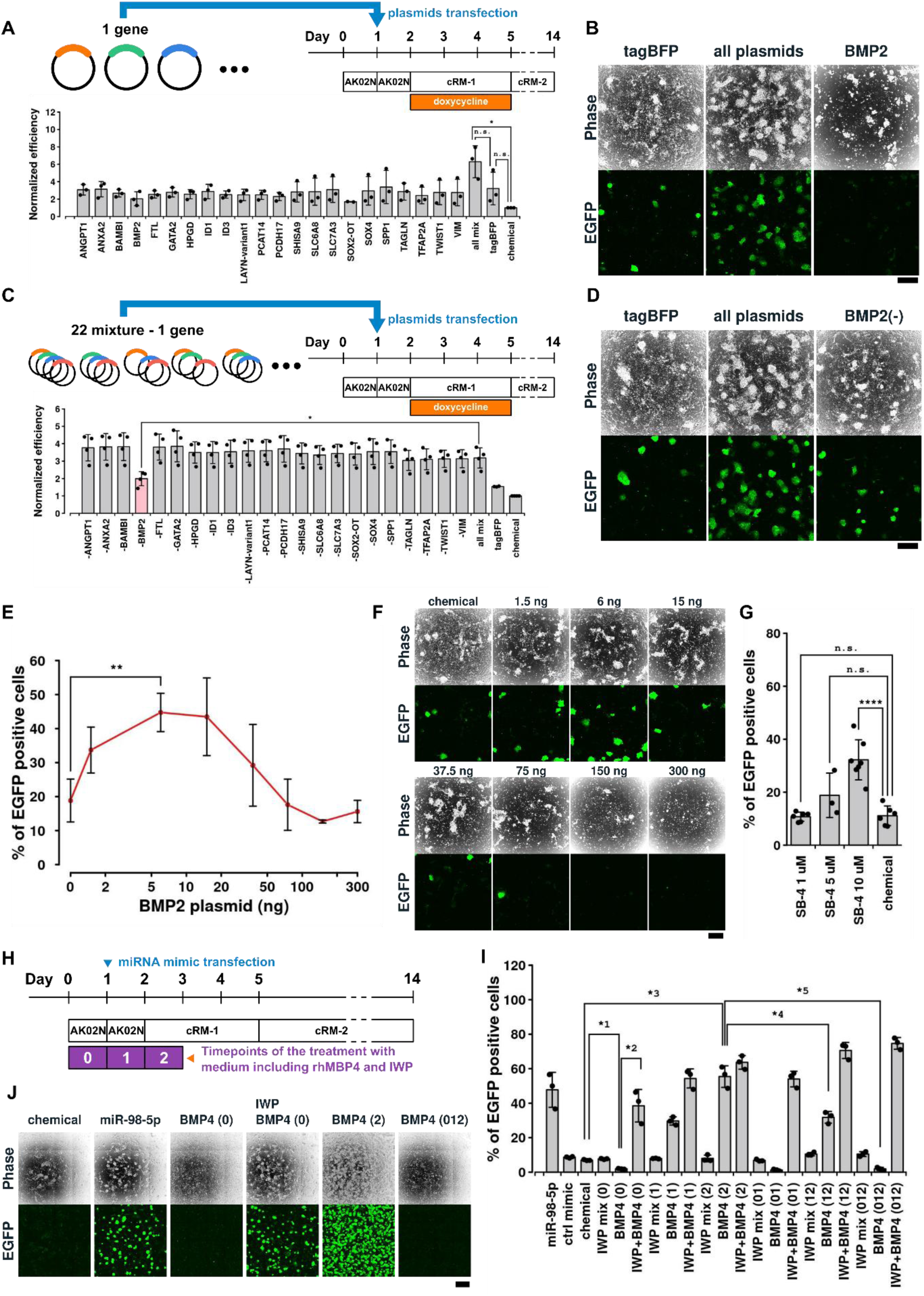
Effects of upregulated candidate genes on resetting efficiency. A: Resetting efficiency of cells transfected with Dox-inducible plasmids expressing candidate genes in the presence of doxycycline. The combined plasmid mixture (all mix) enhanced resetting efficiency. Biological replicate (n=3). \**P* = 3.8×10^-2^. n.s. indicates not significant (p-value > 0.05). B: Representative microscopic images of cells in **A** on day 14. Additional images are shown in Figure S4. Scale bar, 500 μm. C: Resetting efficiency of the plasmid mixture without one candidate gene. Removal of the *BMP2* plasmid abolished the enhancement of resetting efficiency. Biological replicate (n>=3). \**P* = 1.80×10^-2.^ D: Representative microscopic images of cells in **C** on day 14. Additional images are shown in Figure S5. Scale bar, 500 μm. E: Effects on the resetting efficiency by varying amounts of the *BMP2* plasmid. (n>=3). The x-axis is log-scaled for better data visualization. \*\**P* = 9.36×10^-3^. F: Microscopic images of cells in **E** on day 14. Scale bar, 500 μm. G: Resetting efficiency of cells treated with BMP signaling activator, SB-4, on day 14. \*\*\*\**P* = 1.11×10^-4^. n.s. indicates not significant (p-value > 0.5). H: Resetting cells using a medium with rhBMP4 and/or WNT inhibitor cocktail (IWP cocktail). I: Resetting efficiency of cells treated with rhBMP4 and/or IWP cocktail. Numbers in parentheses (0, 1, or 2) represent the duration (days) of treatment with rhBMP4 and/or IWP cocktail in **H**. Biological replicate (n=3). *1 *P* = 5.64×10^-5^. *2 *P* = 2.14×10^-2^. *3 *P* = 5.43×10^-3^. *4 *P* = 5.79×10^-3^. *5 *P* = 5.16×10^-3^. J: Representative images of cells treated with rhBMP4 and/or IWP cocktail on day 14. Scale bar, 1 mm.

To determine the contribution of each gene, we systematically omitted one gene at a time from the 22-plasmid mixture. Exclusion of *BMP2* markedly reduced the proportion of EGFP-positive cells compared with the complete mixture (Figures 4C, D, S5), indicating a key role for BMP2. However, *BMP2* plasmid transfection alone did not enhance resetting (Figure 4A). Notably, many BMP2-transfected cells eventually underwent cell death (Figure 4A, B, S6A). Titration experiments revealed that resetting efficiency peaked when ∼6–15 ng of BMP2 plasmid was used per 24-well plate (1 × 10⁵ cells/well) (Figure 4E). Higher plasmid doses caused cell death (Figure 4F), which occurred not immediately after transfection but rapidly from day 7 onward (Figure S6A). These results suggest that excessive BMP2 activity above a certain threshold, rather than plasmid toxicity, accounts for the observed cell death. Consistently, treatment with the BMP activator SB-4 increased the number of EGFP-positive cells (Figure 4G), confirming that BMP signaling promotes resetting.

BMP signaling has previously been shown to drive differentiation of iPSCs/ESCs into mesodermal lineages (e.g., mesenchymal stem cells, chondrocytes, cardiomyocytes),^40,41^ seemingly at odds with its role in naïve induction, which is associated with a more embryonic state. Our findings, however, indicate that BMP signaling can promote resetting to the naïve state. We therefore infer that BMP signaling enhances resetting only within a restricted temporal window—analogous to the effect of miR-98-5p, which improved resetting when applied on day 1 but not on day 2 (Figure S1B).

Accordingly, we added human recombinant BMP4 (rhBMP4) to the culture medium at different time points to test whether BMP signaling contributes to resetting. A significant increase in EGFP-positive cells was observed when rhBMP4 was added at the time of medium replacement with cRM-1 (Figure 4H–J). Conversely, the BMP inhibitor, LDN193189, abolished the enhancement of resetting efficiency induced by miR-98-5p (Figure S6B), indicating a strong involvement of BMP signaling in this process. The effect of rhBMP4 was time-dependent: resetting efficiency improved when rhBMP4 was added at the cRM-1 exchange but decreased when added earlier (Figure 4H-J; rhBMP4(0), rhBMP4(01), and rhBMP4(012); parentheses denote the time points of rhBMP4 or IWP-cocktail treatment).

Because WNT signaling antagonizes BMP activity and our scRNA-seq analysis showed that WNT1 expression was suppressed in miR-98-5p–treated cells at day 8 (Figure 3G), we next examined the role of WNT signaling. Although the WNT inhibitor (IWP-cocktail) alone did not promote resetting, it rescued the reduction in efficiency caused by early rhBMP4 treatment (day 0) (Figure 4I; IWP+BMP4(0) vs. BMP4(0)). WNT signaling has been reported to drive loss of pluripotency and mesodermal differentiation through the BMP4 pathway,^42^ suggesting that inhibiting WNT prevents BMP4-induced differentiation into alternative lineages.

Finally, we compared rhBMP2 and rhBMP4 and found no significant difference (Figure S6C), implying that BMP2/4 signaling in general, rather than rhBMP4 specifically, facilitates resetting. Together, these results demonstrate that transient, dose- and time-dependent activation of BMP signaling by miR-98-5p promotes resetting and highlights the existence of a critical temporal window for BMP activity.

### NR6A1 downregulation by miR-98-5p contributes to primed-to-naïve conversion

scRNA-seq analysis revealed that several genes, including NR6A1, were downregulated in miR-98-5p mimic–treated cells on day 3 (Figure 3G). Because TargetScan predicts some of the downregulated genes including NR6A1 as direct targets of miR-98-5p,^43^ we hypothesized that its repression plays a central role in the resetting process. To test this, we performed siRNA knockdown experiments on 20 candidate genes reduced by miR-98-5p in Cluster 0. Among these, siRNA against NR6A1 (siNR6A1) significantly increased the number of EGFP-positive cells (Figure 5A), suggesting that NR6A1 suppresses the conversion of primed ESCs to naïve PSCs. This is consistent with previous studies showing that NR6A1 blocks *NANOG* expression in ESCs and is linked to pluripotency regulation ^44^.

**Figure 5.**
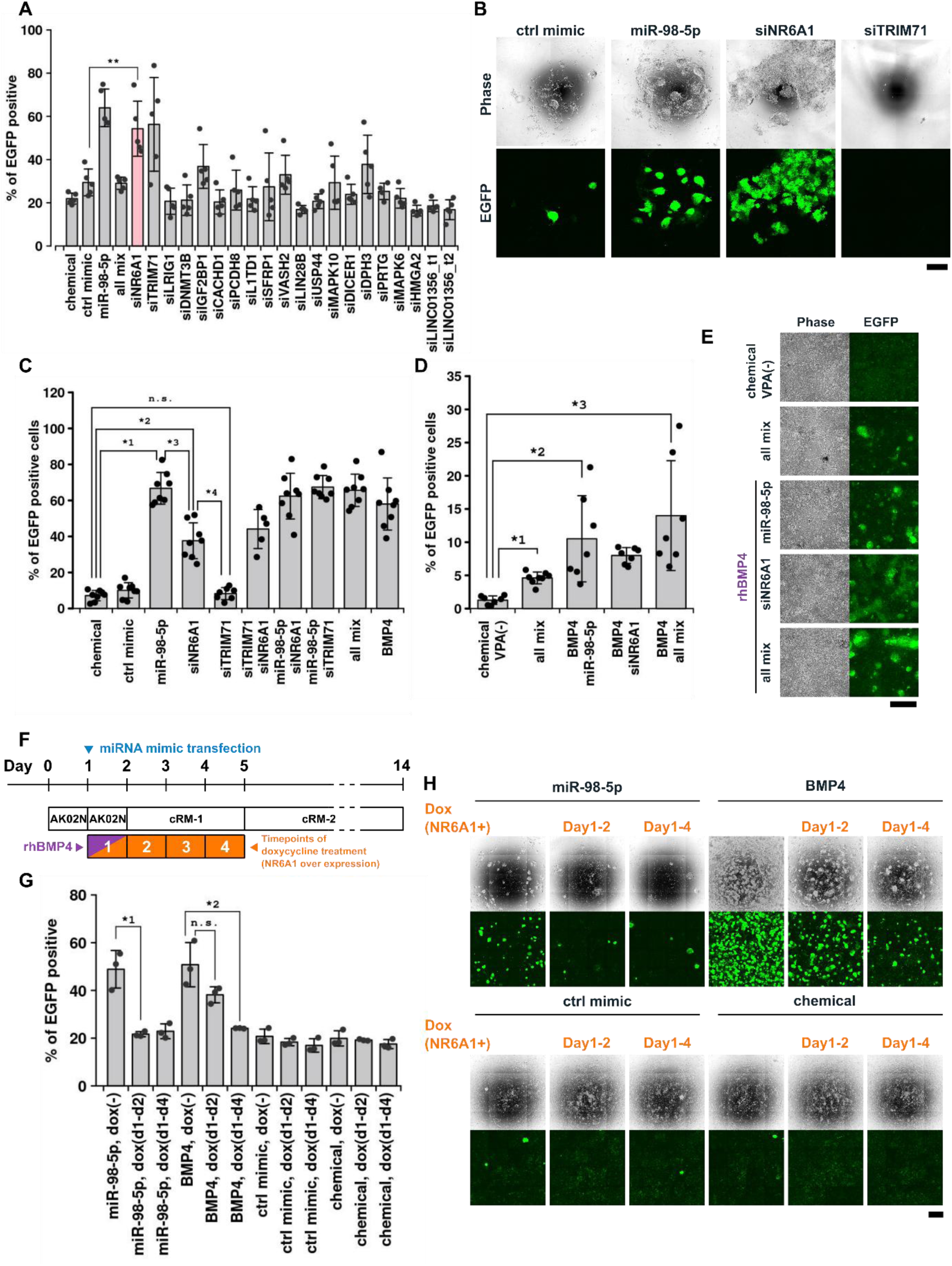
Effects of downregulated candidate genes on resetting efficiency. A: Candidate gene screening using a 96-well plate format siRNA library. siLINC01356_t1 and siLINC01356_t2 are siRNAs targeting different regions in LINC01356. Other wells contained mixtures of four siRNAs per gene. Biological replicate (n=3). \*\**P* = 8.28×10^-3^. B: Microscopic images of cells in **A** on day 14. Cells treated with siTRIM71 were almost entirely lost. Scale bar, 500 μm. C: Resetting efficiency of the combination of siNR6A1, siTRIM71, miR-98-5p mimic, and rhBMP4. “all mix” refers to the mixture of miR-98-5p mimic, siNR6A1, and siTRIM71. Biological replicate (n>=5). *1 *P* = 4.2×10^-5^. *2 *P* = 2.9×10^-5^. *3 *P* = 2.5×10^-5^. *4 *P* = 3.0×10^-5^. n.s., not significant. D: Resetting efficiency of the combination of siRNAs, miR-98-5p mimic, and rhBMP4 without VPA. “all mix” refers to the the mixture of miR-98-5p mimic, siNR6A1, and siTRIM71. *1 *P* = 1.54×10^-6^. *2 *P* = 9.13×10^-3^. *3 *P* = 6.48×10^-3^. E: Microscopic images of cells in **D** on day 14. “all mix” refers to the mixture of miR-98-5p mimic, siNR6A1, and siTRIM71. Scale bar, 500 μm. F: Schematic of resetting inhibition by doxycycline-induced NR6A1 overexpression. G: Effects of NR6A1 overexpression on resetting efficiency. miR-98-5p or ctrl mimics were transfected into a Dox-inducible NR6A1-overexpressing H9EOS cell line, or rhBMP4 was added to the culture medium. EGFP-positive cells were measured on day 14. “d1–d2” and “d1–d4” denote periods of doxycycline treatment (from day 1 to day 2 and from day 1 to day 4, respectively). *1 *P* = 2.50×10^-2^. *2 *P* = 3.80×10^-2^. n.s., not significant. H: Microscopic images of cells shown in **F**, **G** on day 14. Scale bar, 1 mm.

TargetScan also predicts TRIM71 as a target of miR-98-5p.^43^ However, siRNA against TRIM71 (siTRIM71) had only modest effects, and did not increase EGFP-positive cells significantly (Figure 5B), likely due to promoting cell death in the presence of VPA (Figure S7A). TRIM71 has been reported to suppress apoptosis via ubiquitin-dependent degradation of p53,^45^ which may relate to the reduction of non-naïve-like cells observed after miR-98-5p treatment (Figure 1H). Thus, contribution of TRIM71 to resetting appears secondary compared with NR6A1.

We next scaled up the experiments using a 48-well format to assess combinatorial effects of miR-98-5p, rhBMP4, siNR6A1, and siTRIM71. While siTRIM71 alone did not increase resetting efficiency, siNR6A1 alone enhanced resetting, though less effectively than the miR-98-5p mimic (Figure 5C). Moreover, EGFP-positive cells appeared earlier in siNR6A1-treated cultures than in miR-98-5p–treated cultures (Figure S7B). By day 14, both treatments produced EGFP-positive colonies, but siNR6A1 colonies were flat, whereas miR-98-5p colonies exhibited dome-shaped morphology. The difference in morphology implies that the effect of miR-98-5p and siNR6A1 overlap but are not identical. Importantly, non-naïve (EGFP-negative) cells were more effectively reduced by miR-98-5p, likely due to additional gene regulatory effects beyond NR6A1 knockdown (Figures 1H, 3A).

Combining miR-98-5p, rhBMP4, and siNR6A1 further increased resetting efficiency, even without the HDAC inhibitor VPA (Figures 5D, E; Figure S7A), suggesting that miR-98-5p and BMP signaling can bypass the broad chromatin remodeling usually required for resetting.

To further confirm NR6A1’s role, we generated a Dox-inducible NR6A1-overexpressing H9EOS line. Overexpression of NR6A1 during days 1–2 or 1–4 significantly suppressed resetting induced by both miR-98-5p and rhBMP4 (Figures 5F–H), establishing that NR6A1 is a key inhibitory factor directly targeted by miR-98-5p.

### RNA-seq analysis reveals cell type changes and rhBMP4-mediated effects

We next assessed the similarity of naïve-like hPSCs generated by rhBMP4 or miR-98-5p to previously reported naïve hPSCs. Naïve-like hPSCs were induced from H9EOS cells with rhBMP4 or miR-98-5p, and EGFP-positive cells were sorted on day 28 for RNA-seq analysis. At this stage, cells treated with either miR-98-5p or rhBMP4 expressed key naïve markers, including *DNMT3L, NANOG, TBX3, SUSD2, DPPA3, and KLF17* (Figure 6A). When compared with previously published RNA-seq data (GSE144994),^6^ our primed hPSCs clustered closely with database primed samples, while our day 28 naïve-like hPSCs clustered closely with reported naïve hPSCs (Figure 6B). Consistently, principal component analysis positioned our day 28 naïve-like hPSCs near published naïve hPSCs (Figure 6C), demonstrating strong transcriptional similarity. Notably, although BMP4 markedly enhanced resetting efficiency, the transcriptional differences between BMP4-treated and untreated cells were less pronounced than those observed between VPA-treated and untreated cells during the early stages of resetting.

**Figure 6.**
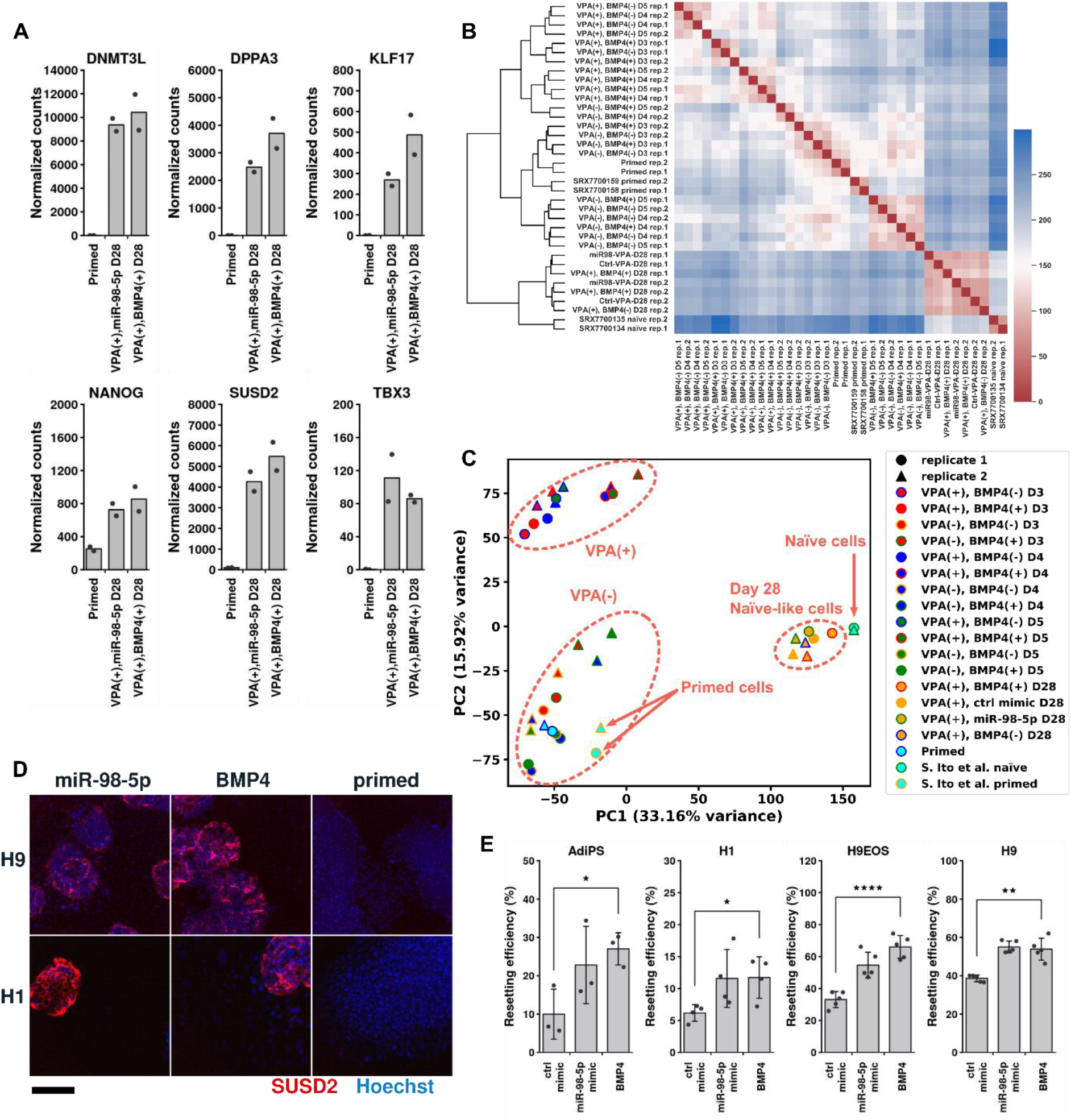
Transcriptomic and phenotypic characterization of naïve-like hPSCs generated by miR-98-5p/rhBMP4. A: Normalized read counts of naïve markers (DNMT3L, DPPA3, KLF17, NANOG, SUSD2, TBX3) in primed hPSCs and EGFP-positive cells after miR-98-5p/rhBMP4 treatment on day 14. B, C: Distance heatmap with hierarchical clustering (Ward method) and PCA plot of gene expression profiles in whole cells on days 3, 4, and 5, and in EGFP-positive cells on day 28 with or without VPA and rhBMP4. “miR-98-5p” indicates cells transfected on day 1. Naïve hPSC reference datasets were obtained from NCBI GEO (GSE144994). D: Representative immunofluorescent staining of SUSD2 in H9 and H1 cells on day 14 after miR-98-5p/rhBMP4 treatment or in primed culture. Scale bar, 200 μm. E: Percentage of SUSD2-positive cells in different hPSC lines (AdiPS, H1, H9EOS, H9) on day 14 after miR-98-5p/rhBMP4 treatment. Biological replicate (n=3). P-values from left to right: \**P* = 2.54×10^-2^, \**P* = 3.47×10^-2^, \*\*\*\**P* = 5.43×10^-5^, and \*\**P* = 2.76×10^-3^.

Antibody staining of the naïve hPSC surface marker SUSD2 showed expression that correlated with the increase in EGFP-positive cells (Figure S8). Increased SUSD2 expression was also observed across multiple hiPSC and hESC lines, including AdiPS, H1, and H9 cells, after resetting with either miR-98-5p or rhBMP4 (Figure 6D, E; Figure S8). Immunostaining further revealed that SUSD2 expression was associated with dome-shaped colony morphology, a hallmark of naïve hPSCs (Figure 6D, Figure S8).

Together, these findings indicate that resetting with miR-98-5p or rhBMP4 produces naïve-like hPSCs that closely resemble reported naïve hPSCs in terms of gene expression, surface marker profiles, and morphology. These results suggest that miR-98-5p and BMP4 promote resetting through mechanistically convergent pathways, with miR-98-5p engaging BMP signaling, and that both ultimately drive phenotypic convergence toward the naïve state.

## Discussion

Our study reveals that a let-7 family miRNA, miR-98-5p, enhances the transition in cell fate from human primed PSCs (ESCs or iPSCs) to the naïve state. To the best of our knowledge, this is the first demonstration that miRNA can facilitate the generation of naïve hPSCs. Notably, we discovered that transient activation of BMP2/4, upregulated two days after miR-98-5p transfection, greatly facilitates naïve hPSC resetting (Figure 4I, J).

Interestingly, the miR-98-5p and let-7 family miRNAs dramatically enhanced the resetting efficiency, likely by regulating the transient expression of multiple genes. scRNA-seq analysis revealed that signaling genes involved in differentiation, including *BMP2, WNT1, and NR6A1,* are affected by miR-98-5p mimic. Among them, rhBMP4 directly induced resetting, whereas NR6A1 overexpression suppressed it. In addition, both miR-98-5p and rhBMP4 act only during a limited time window during resetting and do not promote resetting otherwise. Specifically, rhBMP4 addition before day 2 significantly impaired resetting efficiency (Figure 4H, I, J), consistent with its well-established role in promoting mesoderm and endoderm differentiation.^46^ Thus, adding rhBMP4 prior to VPA treatment may trigger lineage commitment, leading to loss of pluripotency.

We found that *NR6A1* and *TRIM71* are downregulated by miR-98-5p, and both are predicted miR-98-5p targets by TargetScan.^43^ NR6A1 suppresses primed-to-naïve conversion and antagonizes the resetting promoted by miR-98-5p and rhBMP4 (Figure 5F–H). NR6A1 is involved in germ line formation^47^ and neural lineages^48^ and suppresses genes that maintain pluripotency, such as Sox2, Nanog, and Oct4.^44,49^ Therefore, the expression of *NR6A1* may promote aspects of germline development, and, in turn, decrease the resetting efficiency (Figure 7A). In contrast, TRIM71 is known to suppress apoptosis through p53 degradation.^45^ Consistent with this, our results indicate that TRIM71 may help eliminate cells that fail to acquire a naïve-like state, suggesting that miR-98-5p enhances resetting efficiency not only by promoting naïve induction (Figure 2E) but also by facilitating the selective removal of inappropriate cell states (Figures 1H).

**Figure 7.**
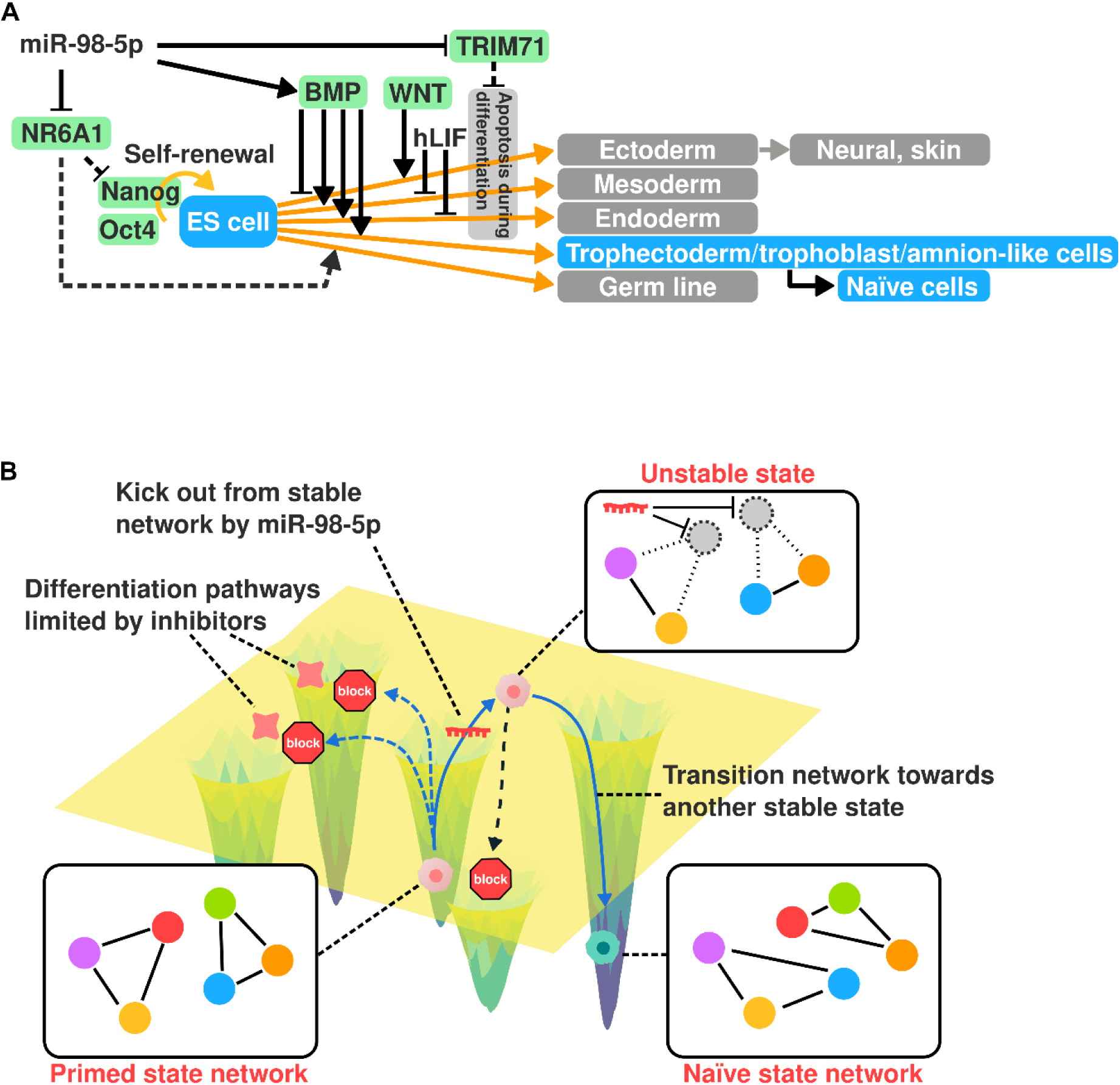
Model of miRNA-driven primed-to-naïve resetting. A: Proposed mechanism by which miR-98-5p promotes resetting. miR-98-5p directly suppresses *NR6A1* and *TRIM71*. Downregulation of *NR6A1* relieves repression of the pluripotency factors *NANOG* and *OCT4* and may prevent premature differentiation. NR6A1 has also been implicated in later stages of germline development and maintenance. *TRIM71* suppression prevents p53-mediated apoptosis in pluripotent cells during differentiation and may also facilitate the elimination of non-naïve differentiated cells. In addition, hLIF restricts mesoderm and endoderm differentiation, while promoting trophectoderm and amnion-like cell fates via activation of BMP signaling. B: Conceptual model of miRNA-driven resetting from the primed to the naïve state. Primed cells reside in a stable attractor and cannot spontaneously transition. miRNA perturbs this stable gene-expression network by simultaneously downregulating multiple targets, providing the drive to escape into an intermediate unstable state. The network then stabilizes into the naïve state, as differentiation pathways are constrained by various inhibitors.

Cell type prediction with ScType indicated that miR-98-5p-transfected cells may transiently pass through trophectoderm-or amnion-like states during the transition to naïve hPSCs (Figure 7A). This trajectory is consistent with previous studies reporting that human and monkey naïve hPSCs share features with trophoblast or amnion cells,^6,39,50,51^ and with the report by Guo and colleagues suggesting that naïve and primed hPSCs exhibit lineage potentials toward trophectoderm and amnion, respectively, thereby linking pluripotency states to extraembryonic trajectories.^38^ The observation that miR-98-5p enhances BMP2 expression and that cells with trophoblast-like signatures emerge during resetting (Figure S3G, I) is consistent with prior findings that BMP4 induces trophoblast stem cells. Furthermore, since mouse naïve PSCs can arise via PGC-like intermediates,^52^ our identification of NR6A1 (a germline-associated factor) as a barrier further supports the notion that naïve hPSCs occupy a state proximal to germline, trophectoderm, and amnion lineages.

Although BMP4 and NR6A1 are generally regarded as unidirectional regulators of differentiation,^34,40,47,51^ we found that they can also play important roles in the resetting process in the opposite direction of differentiation. This is analogous to our unexpected observation that miR-98-5p, a member of the let-7 family traditionally associated with promoting differentiation, instead facilitates naïve-state induction. These findings suggest that a key function of miR-98-5p may be to destabilize the gene network of hESCs/iPSCs and enhance cellular plasticity by shifting the established, stable network from a ground state to an excited state, rather than simply promoting differentiation or resetting in a linear manner. This aligns with the broader principle in reprogramming that an initial “network destabilization” or stochastic phase often precedes deterministic cell fate conversion.^53^ Typically, differentiation into ectoderm, mesoderm, and endoderm is inhibited by BMPs and hLIF. Furthermore, NR6A1 is normally silenced, thereby restricting maturation of germline and neural lineages.^47,48^ As a result, by suppressing differentiation signals during the transition from the primed to the naïve state, cells are permitted to shift toward the naïve state (Figure 7A, B).

Here, through screening with a miRNA-responsive mRNA library,^56–58^ we discovered that miR-98-5p facilitates resetting to naïve hPSCs. Introducing a synthetic miR-98-5p mimic during the initial stages of the naïve resetting protocol significantly enhanced induction efficiency (Figures 1, 2). However, quantitative miRNA expression analysis (AQ-Seq)^29^ revealed no substantial difference in miR-98-5p levels between naïve hPSCs and primed hESCs. Indeed, previous studies have reported conflicting results regarding miR-98-5p expression in naïve hPSCs, with some suggesting high expression^24^ and others low expression.^25^ Our own miRNA expression analyses in naïve hPSCs, purified by removing MEFs using gelatin-coated wells, showed a tendency toward higher miR-98-5p levels, suggesting that earlier studies may have been confounded by MEF-derived contamination of let-7 family miRNAs, including miR-98-5p. In addition, transient upregulation of let-7 family members, including miR-98-5p, has been reported during mouse iPSC induction.^18^

Taken together, these findings suggest the following possibilities for the role of miR-98-5p in naïve induction: its expression and activity are not necessarily high in naïve hPSCs (and may fluctuate during resetting), but its ectopic introduction into primed ESCs destabilizes pre-existing stable gene networks. This destabilization may directly or indirectly regulate key genes such as BMP2 and NR6A1, thereby enhancing resetting efficiency. By contrast, conventional iPSC reprogramming via transcription factors (Yamanaka factors) or miRNAs (e.g., miR-302 in hPSCs^19^) has relied on the forced expression of factors that are already highly expressed in the target state. Intriguingly, even let-7 miRNAs, which are not highly expressed in naïve hPSCs, can promote naïve induction by perturbing the stability of the initial primed network. This highlights a conceptually distinct strategy: rather than enforcing the expression of lineage-defining factors, reprogramming can also be achieved by network destabilization of the starting cells through transient perturbations. Such an approach, based on disrupting established gene regulatory circuits, may be broadly applicable to other cell fate conversion strategies and the generation of alternative target cell types.

Our results revealed that forced and transient expression of miR-98-5p promotes the resetting process. However, this effect is likely specific to artificial reprogramming and does not necessarily imply that miR-98-5p is transiently expressed during natural development. One possible explanation is that evolutionarily conserved let-7 family–responsive sequences are widely embedded in genes forming the primed ES/iPSC network, making these networks particularly susceptible to perturbation by synthetic

miR-98-5p. Whether miR-98-5p or other let-7 family miRNAs are naturally involved in naïve conversion remains unclear from our study. Nevertheless, previous reports have shown that naïve-like conversion of primed hPSCs can be facilitated by RNA transfer from surrounding cells during coculture.^59^ Since feeder cells often express high levels of let-7 family miRNAs, it is conceivable that these miRNAs may be supplied externally. Indeed, naïve induction methods using feeder cells are reported more frequently than feeder-free approaches, supporting the potential involvement of exogenous let-7 input. Thus, while our findings demonstrate that synthetic miR-98-5p can efficiently enhance resetting, the observed effects may reflect an artificial reprogramming mechanism rather than a naturally occurring developmental process, highlighting the need for further detailed investigation.

In conclusion, this study provides the first evidence that a let-7 family miRNA, miR-98-5p, can promote primed-to-naïve state conversion. By demonstrating that transient, synthetic delivery of miR-98-5p efficiently enhances resetting, we establish a miRNA-based strategy for generating naïve hPSCs. This approach not only uncovers a previously unrecognized role for let-7 family miRNAs in destabilizing pluripotency networks but also offers a feeder-free, genome-safe platform with strong potential for regenerative medicine. These findings expand our understanding of human pluripotency and highlight synthetic miRNAs and transient BMP 2/4 activation as a promising tool for both mechanistic studies and clinical translation. More broadly, our results suggest that miRNA-induced network destabilization may represent a generalizable principle for facilitating cell fate transitions, providing a conceptual framework that could be leveraged across diverse reprogramming and differentiation contexts.

## Resource Availability

Requests for further information and resources should be directed to and will be fulfilled by the lead contact, Yoshihiko Fujita or Hirohide Saito. This study did not generate new unique reagents. All plasmids, synthetic RNA constructs, and naïve hPSC lines established in this work are available from the lead contact upon reasonable request and completion of a material transfer agreement.

## Acknowledgments

We thank Dr. Narry Kim (Seoul National University) for providing information and valuable advice on the AQ-Seq protocol, and Dr. Kelvin K. Hui (Kyoto University) for proofreading the manuscript. We also thank Dr. Michinori Saitou (Kyoto University), Dr. Konrad Hochedlinger (Harvard University) for helpful discussions, and Hiromi Takemoto for their administrative support. This work was supported by the JSPS KAKENHI (Grant Numbers JP20H05626, JP21K06199), AMED (Grant Number JP21bm0704032h0003), and iPS Cell Research Fund from Center for iPS Cell Research and Application (Grant Number JP23bm1323001h0001), Kyoto University.

## Author contribution

Y.F. and H.S. conceived and designed the project. Y.F., M.H., and K.H. performed the experiments. Y.F. and T.Y. analyzed the data. B.K. and M.U. prepared naïve cells by the feeder protocol for screening and RNA-seq. H.O. prepared miRNA-seq library. Y.F., M.H., and H.S. wrote the manuscript. M.L., T.Y., and Y.T. discussed the results and commented on the manuscript.

## Data availability

All data needed to evaluate the conclusions in the paper are present in the paper and/or the Supplementary Materials. All RNA-seq data generated in this publication have been deposited in NCBI’s Gene Expression Omnibus (Edgar et al., 2002)^60^ and are accessible through GEO Series accession number GSE280065 (https://www.ncbi.nlm.nih.gov/geo/query/acc.cgi?acc=GSE280065).

## Declaration of generative AI and AI-assisted technologies

During the preparation of this work, we used ChatGPT for proofreading. After using this service, we reviewed and edited the content as needed and took full responsibility for the content of the published article.

## Inclusion & Ethics

This study was conducted in accordance with the ethical guidelines and policies approved by the CiRA (Center for iPS Cell Research and Application) Ethics Committee. The research plan was reviewed and approved by the committee to ensure compliance with ethical standards and proper handling of all materials and data.

## Methods

### Cell culture and chemical resetting

Pluripotent stem cells (H9-NK, H9EOS, H9EOS-NR6A1, H1, 1383D4, and AdiPS) were cultured in AK02N (Ajinomoto, AK02N) medium without feeder cells at 37°C with 5% CO_2_ and were passaged approximately every week. Cells were washed with PBS and incubated with Accutase (Nacalai Tesque, 12679-54) at 37°C for at least 10 min before pipetting. After pipetting, cells were centrifuged at 160 × *g* for 5 min, and the supernatant was removed by aspiration. Cells were resuspended in AK02N medium containing 10 μM Y-27632 (Wako, 036-24023) and 1.6 ng/μL iMatrix-511 silk (Nippi, 387-10131) and transferred to a new plate or were resuspended in AK02N medium containing 10 μM Y-27632 and transferred to a new plate pre-coated by iMatrix-511 silk.

For chemical resetting, cells were seeded in a multi-well plate in AK02N containing 10 μM Y-27632 and 1.6 ng/μL iMatrix-511 silk and replaced on day 1 with AK02N. From day 2 to day 5, cells were cultured in cRM-1 medium (NDiff 227 (TaKaRa, Y40002) medium containing 1 μM PD0325901, 10 ng/mL human LIF (hLIF), and 1 mM valproic acid (VPA)). From day 5, cells were cultured in cRM-2 medium (NDIff 227 medium containing 1 μM PD0325901, 10 ng/mL hLIF, 2 μM Go6983 (R&D, 2285), and 20 μM XAV939 (Sigma, X3004-5MG)).

### Flow cytometric analysis

Cells were harvested by AccuMax, followed by incubation at 37°C for about 10 min. After confirming cell detachment from well bottoms, suspended cells were analyzed using an Accuri C6 flow cytometer (BD Bioscience) with the FL1 filter (530/30 nm) for EGFP. The data were extracted from FCS files generated by the equipment using the “flowCore” package of R^61^ and analyzed using the package “mgcv”. Outliers were removed using the package “outliers”. The FSC/SSC gates were set to enclose the densely populated cell clusters observed in the plots. Since GFP-positive cells also appeared in the cells treated with control miRNA mimic, the exact separation of GFP-positive and GFP-negative cell populations could not be strictly defined. To address this limitation, controls were consistently included, allowing for the relative evaluation of GFP expression.

### miRNA screening

H9-NK or H9EOS cells were seeded in a 12-well plate (6×10^4^ cells/well) and cultured according to the chemical resetting protocol except for transfection of miRNA mimics. 1.6 pmol of miRNA mimics were transfected using Lipofectamine RNAiMAX Transfection Reagent (Thermo Fisher Scientific) according to the manufacturer’s protocol after a medium change on day 1.

### Counting colonies of naïve-like cells

We analyzed 16-bit grayscale fluorescence images in Python (NumPy/SciPy/scikit-image). Images were downsampled (down=8), then rescaled to [0,1]. Background was removed with rolling-ball subtraction (radius 50 pixel). Segmentation used local thresholding (threshold_local, block size 51) with an additive offset, followed by remove_small_objects (min_size = min_area), hole filling, and binary_opening (disk=2). Detected regions (regionprops) were accepted as colonies if area ≥ min_area, mean intensity ≥ min_meanint, and solidity ≥ 0.70. Hyperparameters were chosen by a grid search over gamma, offset, min_meanint, and CLAHE on/off. The single best setting was then applied uniformly to all images.

### Single-cell RNA-seq

H9EOS cells were cultured according to the “Cell culture and chemical resetting” section. Cells were washed by PBS twice and incubated in AccuMax (Nacalai Tesque, 17087-54) at 37°C for more than 15 min. Cells were dissociated by pipetting gently and washed with cold PBS supplemented with 0.04% BSA twice. After filtration by MACS SmartStrainers (30 µm) (Miltenyi Biotec, 130-098-458), single-cell suspensions were loaded into a Chromium Single Cell Controller instrument (10×Genomics) to generate single-cell gel beads in emulsion. 10x Genomics v.3.0 libraries were prepared according to the manufacturer’s instructions (10×Genomics, 1000128). All samples were mixed together and sequenced on an Illumina HiSeq4000 or NovaSeq6000 with paired-end sequencing. scRNA-seq data were mapped and quantified using Cell Ranger 7.1.0 against the hg38 human reference genome (refdata-gex-GRCh38-2020-A, 10x Genomics) including the EOS-EGFP cassette. Raw count data were imported into the Seurat package 4.3.0.^62^ Cells with nfeature > 2,000, nCount between 5,000 and 100,000, and low mitochondrial gene expression (< 15%) were further analyzed. Raw counts were normalized by the NormalizeData() with LogNormalize method and scaled by ScaleData() in the Seurat package. PCA and clustering were performed using the Seurat RunPCA() and FindClusters() with the resolution set at 0.15 (Figure S4-A). Cluster 0 was reanalyzed by FindClusters() with the resolution set at 0.2. Differentially expressed genes in cells transfected with miR-98-5p were identified by FindMarkers(). clusterProfiler 4.8.3^63^ was used for GO analysis. For cell type prediction, we first downloaded human cell markers from CellMarker2.0^36^ and modified the data to adapt to scType software (https://raw.githubusercontent.com/IanevskiAleksandr/sc-type/master/R/sctype_score_.R)^35^. The mean cell type score in each cluster was calculated, and the cell type with the highest score was defined as the predicted cell type in Figure S4-F.

RNA velocity was computed with velocyto (run10x) on each 10x Genomics Cell Ranger sample directory using the appropriate GTF, yielding .loom files with spliced/unspliced counts. For downstream analysis, .loom files from all time points/conditions were loaded into scVelo (v0.3.3). Velocities were inferred with the dynamical model (recover_dynamics → velocity → velocity_graph), followed by latent time estimation and visualization as velocity stream/arrows on UMAP.

### Candidate gene overexpression

Candidate genes were selected from single-cell RNA-seq data with pct1 – pct2 > 0.3 or avg_log2Fold > 0.5. Genes were amplified by RT-PCR from total mRNA and cloned to a doxycycline-inducible vector. Genes that could not be amplified were omitted from the following experiment. For overexpression of each gene, 200 ng of each plasmid and 100 ng of pCAG-PBase were co-transfected into H9EOS cells using Lipofectamine Stem Transfection Reagent according to the manufacturer’s protocol (Thermo Fisher Scientific). For co-expression of all candidate genes except for one gene, the plasmids except for one plasmid were mixed. The 300-ng plasmid mixture and 100 ng of pCAG-PBase were co-transfected into H9EOS cells. Cells were treated with cRM-1 supplemented with 250 ng/mL doxycycline from day 2 to day 4 to express the genes.

### BMP inhibition

H9EOS cells were seeded in laminin-coated 24-well plates in AK02N containing Y-27632 (3×10^4^ cells/well). Before transfection, the medium was replaced with AK02N without Y-27632. 0.8 pmol of miR-98-5p or negative control mimics were transfected using Lipofectamine RNAiMAX Transfection Reagent according to the manufacturer’s protocol. The cells were cultured in the medium supplemented with 100 nM of LDN193189 (STEMGENT, 04-0074) on day 1, day 2, day 3, or day 4. Chemical resetting was conducted according to the section on chemical resetting.

### BMP treatment and WNT inhibition

H9EOS cells were seeded in laminin-coated 24-well plates in AK02N containing Y-27632 (3×10^4^ cells/well). Before transfection, the medium was replaced with AK02N without Y-27632. 0.8 pmol of miR-98-5p or negative control mimics were transfected using Lipofectamine RNAiMAX Transfection Reagent according to the manufacturer’s protocol. Cells were cultured in 10 ng/mL of rhBMP4 or Recombinant Human BMP-2 (rhBMP2) protein (R&D systems) were used. To inhibit WNT signaling, cells were cultured in the presence of 1 µM of IWP cocktail (IWP-2 (Nacalai Tesque, 18175-64), Stemolecule WNT Inhibitor IWP-3 (Reprocell, 04-1935), Stemolecule WNT Inhibitor IWP-4 (Reprocell, 04-0036)). Chemical resetting was conducted according to the section on chemical resetting.

### BMP activator (SB-4) treatment

H9EOS cells were cultured according to the chemical resetting protocol. SB-4 (Tocris, 6881/50) was added during the cRM-1 treatment.

### siRNA screening

Target genes were selected from single-cell RNA-seq data with pct2 – pct1 > 0.2 or avg_log2Fold < - 0.5. H9EOS cells were seeded in a 96-well plate (6×10^3^ cells/well) and cultured according to the chemical resetting protocol except for transfection. After a medium change on day 1, 1 pmol of siRNAs (Dharmacon) and 0.16 pmol of miRNA mimics were transfected using Lipofectamine RNAiMAX Transfection Reagent according to the manufacturer’s protocol.

### *NR6A1* and *TRIM71* knockdown

H9EOS cells were seeded in a multi-well plate and cultured according to the chemical resetting protocol except for transfection. After a medium change on day 1, siRNAs (Dharmacon) and miRNA mimics were transfected using Lipofectamine RNAiMAX Transfection Reagent according to the manufacturer’s protocol. VPA was removed from cRM-1 medium for VPA(-) samples. rhBMP4 was added on day 3 for rhBMP4 (+) samples.

### Establishment of doxycycline-inducible *NR6A1-* or *BFP*-overexpressing H9EOS cells

H9EOS cells were seeded in laminin-coated 24-well plates in AK02N containing Y-27632 (3×10^4^ cells/well). 200 ng pPB-TRE2-NR6A1 -BlaR and 200 ng pCAG-PBase were co-transfected into the cells using Lipofectamine Stem Transfection Reagent according to the manufacturer’s protocol (Thermo Fisher Scientific). Cells were selected using 20 μg/mL of blasticidin (INVIVOGEN 03759-00).

### *NR6A1* overexpression

H9EOS-NR6A1 cells were seeded in laminin-coated 24-well plates in AK02N containing Y-27632 (3×10^4^ cells/well). Before transfection, the medium was replaced with AK02N without Y-27632. 0.8 pmol of miR-98-5p or negative control mimics were transfected using Lipofectamine RNAiMAX Transfection Reagent (Thermo Fisher Scientific) according to the manufacturer’s protocol. Recombinant Human BMP-4 (rhBMP4) protein (Reprocell, 314-BP-010) was added at 10 ng/mL of final concentration on day 2. Cells were cultured in the medium containing 1 μg/mL of doxycycline from day 1 to day 4 or from day 1 to day 2 to overexpress NR6A1. Chemical resetting was conducted according to the section on chemical resetting.

### Fluorescence imaging and image processing

All images were captured by a CQ1 confocal image cytometer (Yokogawa Electric Corporation). Image processing was performed using an ImageJ plugin (https://github.com/yfujita-skgcat/image_converter).

### Plasmid construction

pPB-TRE2-NR6A1_variant1-BlaR *NR6A1* was cloned from H9EOS cell cDNA using the following primers: MHC-595: GACCCTCATTgacctGCCACCatggagcgggacgaaccgc. MHC-596: TGTCGTAACGgacgttcattccttgcccacactggtcttgc. The reaction was conducted using KOD One PCR Master Mix (TOYOBO) with the following conditions: (94°C for 2 min, 40 cycles of 98°C for 10 sec, 55 or 60°C for 5 sec, and 60°C for 13 sec and hold at 15°C). YFP0120-PB-TRE2-PshAl-BlaR was digested by PshAl. These components were assembled using In-Fusion Snap Assembly Master Mix (TaKaRa) according to the manufacturer’s protocol.

### Small RNA-seq

Total RNA was isolated with the mirVana miRNA Isolation Kit (Thermo Fisher Scientific, AM1560/AM1561) according to the manufacturer’s total RNA isolation procedure. Total RNA was eluted with nuclease-free water. RNA concentration was measured with a NanoDrop 2000 spectrophotometer (Thermo Fisher Scientific). The integrity of total RNA samples was measured with an Agilent 2100 Bioanalyzer (Agilent) or a Qsep100 DNA Fragment Analyzer (BiOptic). Libraries for small RNA sequencing were prepared according to the AQ-seq method^29^ with some modifications. Total RNA (2∼5 µg) was resolved on a 15% denaturing polyacrylamide gel, and 17–29 nt small RNA fraction was excised. Excised gel slices were fragmented in 0.5-mL tubes with holes at the bottom by centrifugation at 20,400 × *g* for 10 min to allow the gel to move through the holes to 2-mL tubes. The size-fractioned RNA was eluted from gel slices into 0.3 M NaCl (Nacalai Tesque, 06900-14) at 25°C with overnight shaking at 1,500 rpm. Gel debris was removed with Costar Spin-X centrifuge tubes (Corning, 8162) by centrifugation, and the RNA was ethanol-precipitated with sodium acetate (Nippon Genetics, 316-90081) and GlycoBlue Coprecipitant (Thermo Fisher Scientific, AM9516). The size-fractioned RNA was ligated to 3’ randomized adapter at 25°C for 16 hrs in a reaction mixture containing 0.25 µM 3’ randomized adapter (5’- App/NNNNTGGAATTCTCGGGTGCCAAGG/ddC -3’, purchased from IDT), 200 units of T4 RNA ligase 2 truncated KQ (NEB, M0373), 1× T4 RNA ligase reaction buffer (NEB), 20% PEG 8000 (NEB), and 10 units of SUPERase•In RNase Inhibitor (Thermo Fisher Scientific, AM2694). The adapter-ligated RNA was size-fractioned on 15% denaturing polyacrylamide gel electrophoresis to remove the remaining unligated adapters. The size-fractioned RNA was gel-purified as described above. The adapter-ligated RNA was ligated to 0.18 µM 5’ randomized adapter (5’- GUUCAGAGUUCUACAGUCCGACGAUCNNNN -3’, purchased from IDT) with 14 units of T4 RNA ligase 1 (NEB, M0204), 1X T4 RNA ligase reaction buffer (NEB), 20% PEG 8000 (NEB), 1 mM ATP, and 14 units of SUPERase•In RNase Inhibitor at 37°C for 1 hr followed by heat inactivation of the enzyme at 65°C for 15 min. The adapter-ligated RNA was reverse-transcribed with 200 units of SuperScript III reverse transcriptase (Thermo Fisher Scientific, 18080044), 1× first-strand buffer (Thermo Fisher Scientific), 0.2 µM RT primer (RTP, GCCTTGGCACCCGAGAATTCCA), 0.5 mM dNTP (included in TruSeq Small RNA kit, Illumina), and 5 mM DTT (Thermo Fisher Scientific) at 50°C for 1 hr. Reverse transcriptase was heat-inactivated at 70°C for 15 min after reverse transcription. The cDNA library was amplified with one unit of Phusion High-Fidelity DNA Polymerase (Thermo Fisher Scientific, F530S), 1× Phusion HF buffer (Thermo Fisher Scientific), 0.5 µM primers (RP1 forward primer and RPIX reverse primer from TruSeq Small RNA kit from Illumina), and 0.2 mM dNTP (Takara Bio). The amplified cDNA library was size-fractioned on a 6% non-denaturing polyacrylamide gel to remove adapter dimers. The size-fractioned cDNA library was gel-purified and precipitated as described above. The size of the cDNA library was evaluated with an Agilent 2100 Bioanalyzer or a Qsep100 DNA Fragment Analyzer, with its concentration quantified with KAPA Library Quantification Kits (KAPA Biosystems, KK4873). Pooled cDNA libraries were sequenced on the NextSeq 500 platform (Illumina) with NextSeq 500/550 High Output Kit v2.5 (75 cycles, Illumina).

### RNA sequencing

Cells were cultured basically according to the chemical resetting protocol. For cells without VPA treatment, VPA was removed from cRM-1 medium. For cells treated with rhBMP4, cells were cultured in medium supplemented with 10 ng/mL of rhBMP4 on day 2. For cells transfected with miR-98-5p or control mimics, miRNA mimics were transfected on day 1 after medium change. Cells were harvested on days 3, 4, and 5 after washing with PBS, followed by AccuMax (Nacalai Tesque, 17087-54) treatment.On day 28, GFP-positive cells were sorted using the BD FACSymphony S6 cell sorter. Cell pellets were washed cold PBS once and stored at -80°C until use. RNA was extracted using the mirVana miRNA Isolation Kit (Thermo Fisher Scientific) according to the manufacturer’s protocol. RNA-seq libraries were prepared using the TruSeq Stranded mRNA Library Prep (Illumina) with IDT for Illumina TruSeq RNA UD Indexes (Illumina), quantified using the Qubit dsDNA HS Assay Kit (Thermo Fisher Scientific), and sequenced on a NextSeq500 (Illumina) as 76 bp single reads.

### RNA-seq and AQ-seq data analysis

Sequences obtained by NextSeq500 were trimmed and filtered using fastp 0.23.2.^64^ The human genome, UCSC hg38 August 2015, was downloaded from the Illumina Ready-To-Use Reference Sequences and Annotation website (https://jp.support.illumina.com/sequencing/sequencing_software/igenome.html). Reads were mapped to the customized human genome, including the *EGFP* gene, by STAR 2.7.10b,^65^ and the uniquely mapped reads were counted using htseq-count 2.0.1.^66^ Normalized read counts were calculated using DESeq2 1.40.2 R packages.^67^ The candidate target genes of miR-98-5p were predicted using miRTarBase.^43^ Variance stabilizing transformation for clustering and PCA analysis (Figure 6-B, C) were calculated by DEseq2. The clustering analysis was performed using SciPy 1.10.1 and Seaborn 0.12.2 Python packages. PCA was performed by a Sklearn 1.3.1 Python package. The data of AQ-seq were mapped by miRge3 0.0.9^68^ to human and mouse miRNAs (https://sourceforge.net/projects/mirge3/files/miRge3_Lib/human.tar.gz/download, https://sourceforge.net/projects/mirge3/files/miRge3_Lib/mouse.tar.gz/download).

