## Supplemental files for "Transient let-7/miR-98 reprograms primed pluripotency to induce naive human PSCs"

#### Contents

Supplementary Figure S1. Effects of miRNA on resetting efficiency of H9 cells

Supplementary Figure S2. Genes expression profile on UMAP

Supplementary Figure S3. Assignment of cell type to each cluster in scRNA-seq analysis

Supplementary Figure S4. Effects of overexpression of candidate genes on resetting efficiency

Supplementary Figure S5. Effects of removing candidate genes from gene cocktail on the resetting efficiency

Supplementary Figure S6. Effects of BMP signaling at various times on resetting efficiency

Supplementary Figure S7. Fluorescent images on day 14 after treatment with the combination of siRNA (siNR6A1 and siTRIM71), miR-98-5p, and rhBMP4

Supplementary Figure S8. Fluorescent images of Immunofluorescent staining of SUSD2 of ESCs and iPSCs

#### Supplementary Figure 1

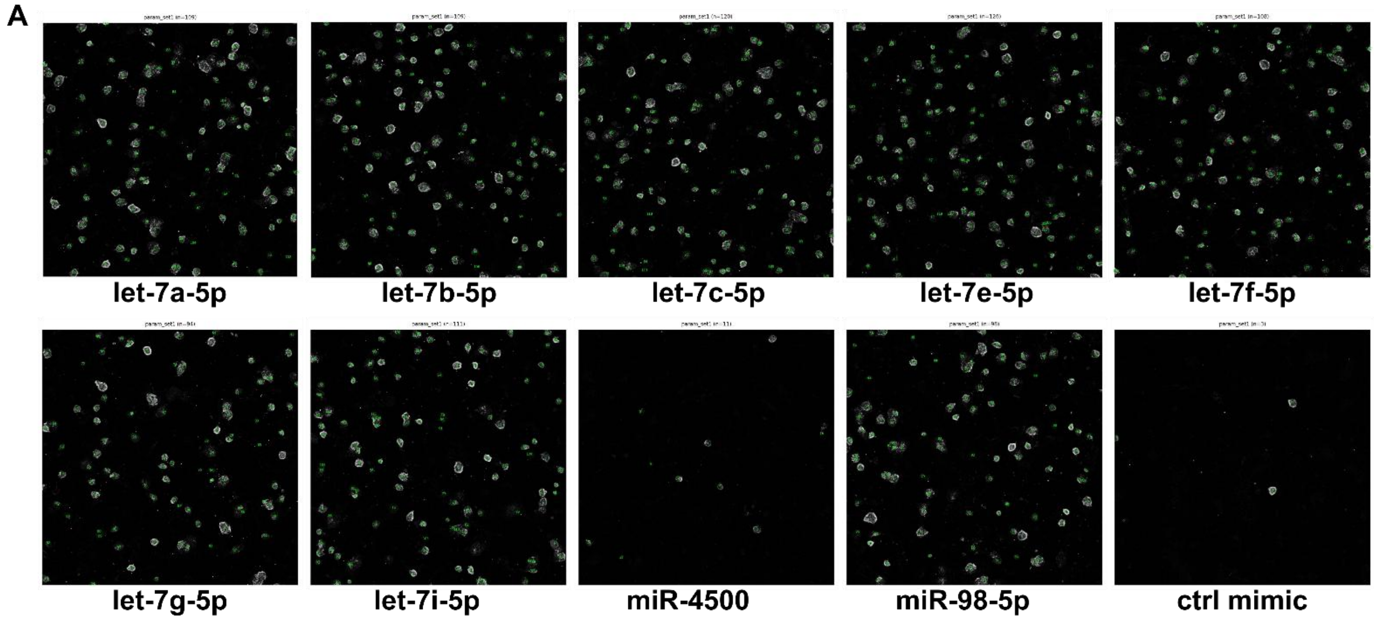


**Supplementary Figure 1. Effects of miRNA on resetting efficiency of H9 cells**

A: Representative images showing colony quantification by image analysis of the images presented in Figure 1D. Green numbers correspond to the serial IDs assigned to each counted naïve-like colony.

### Supplementary Figure 2

#### A naïve marker

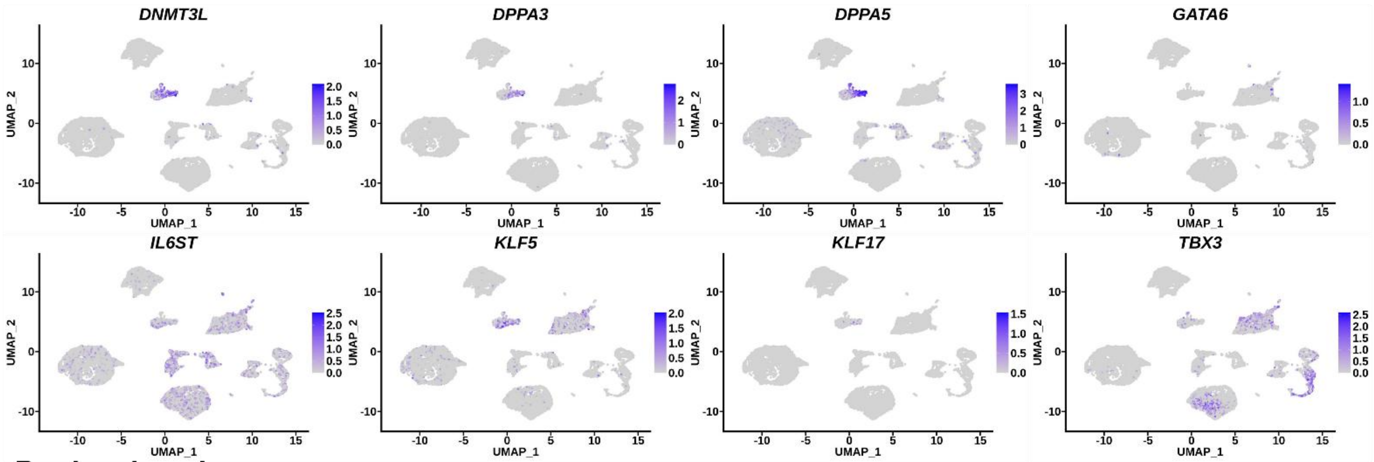


#### B primed marker

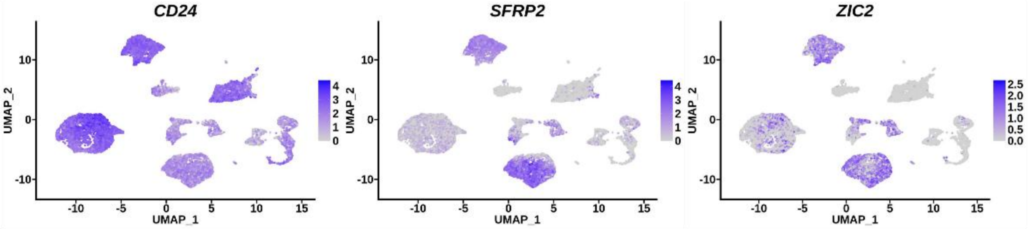


#### C upregulated genes on day 3

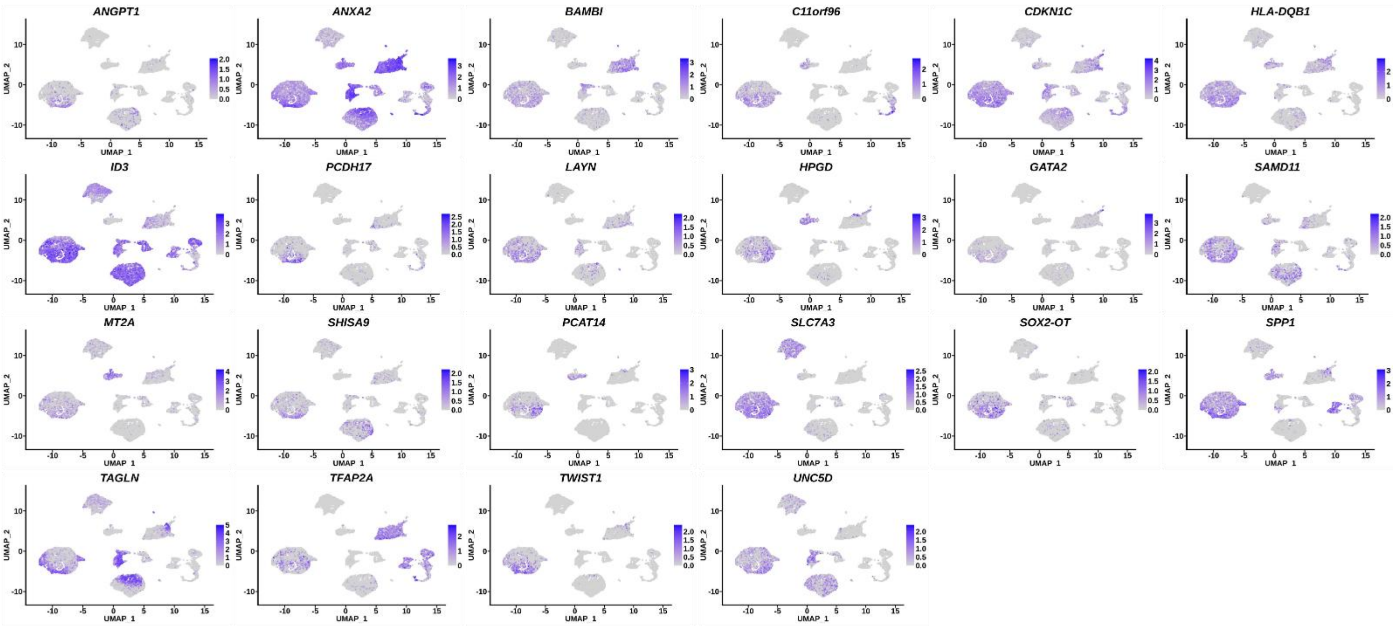


#### D downregulated genes on day 3

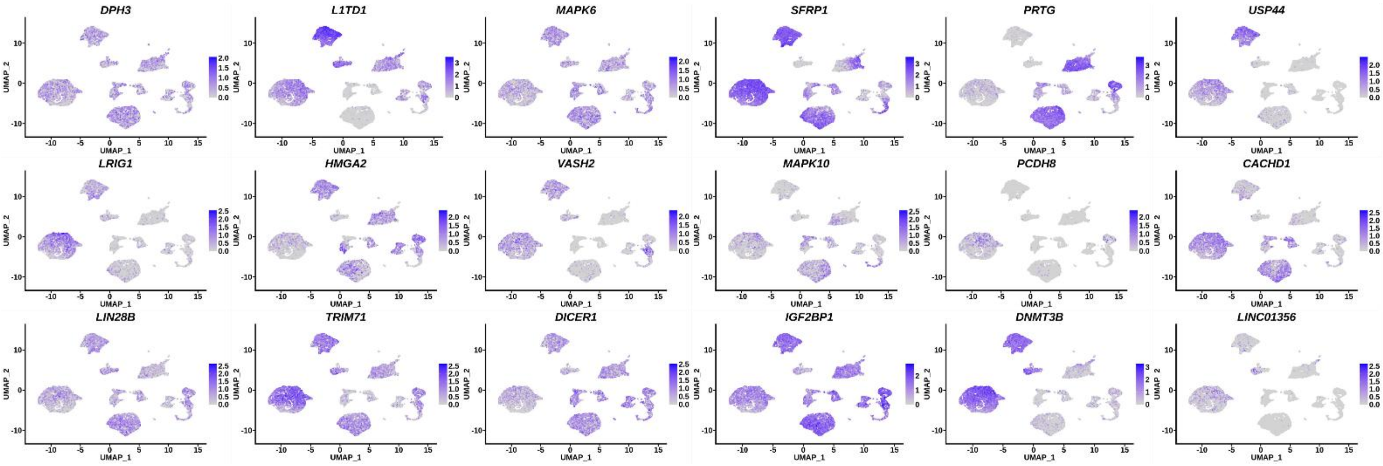

**Supplementary Figure 2. Genes expression profile on UMAP**

A, B: Global expression profiles of naïve and primed markers.

C, D: Examples of up- and downregulated genes in cluster 0 (= day 3).

### Supplementary Figure 3

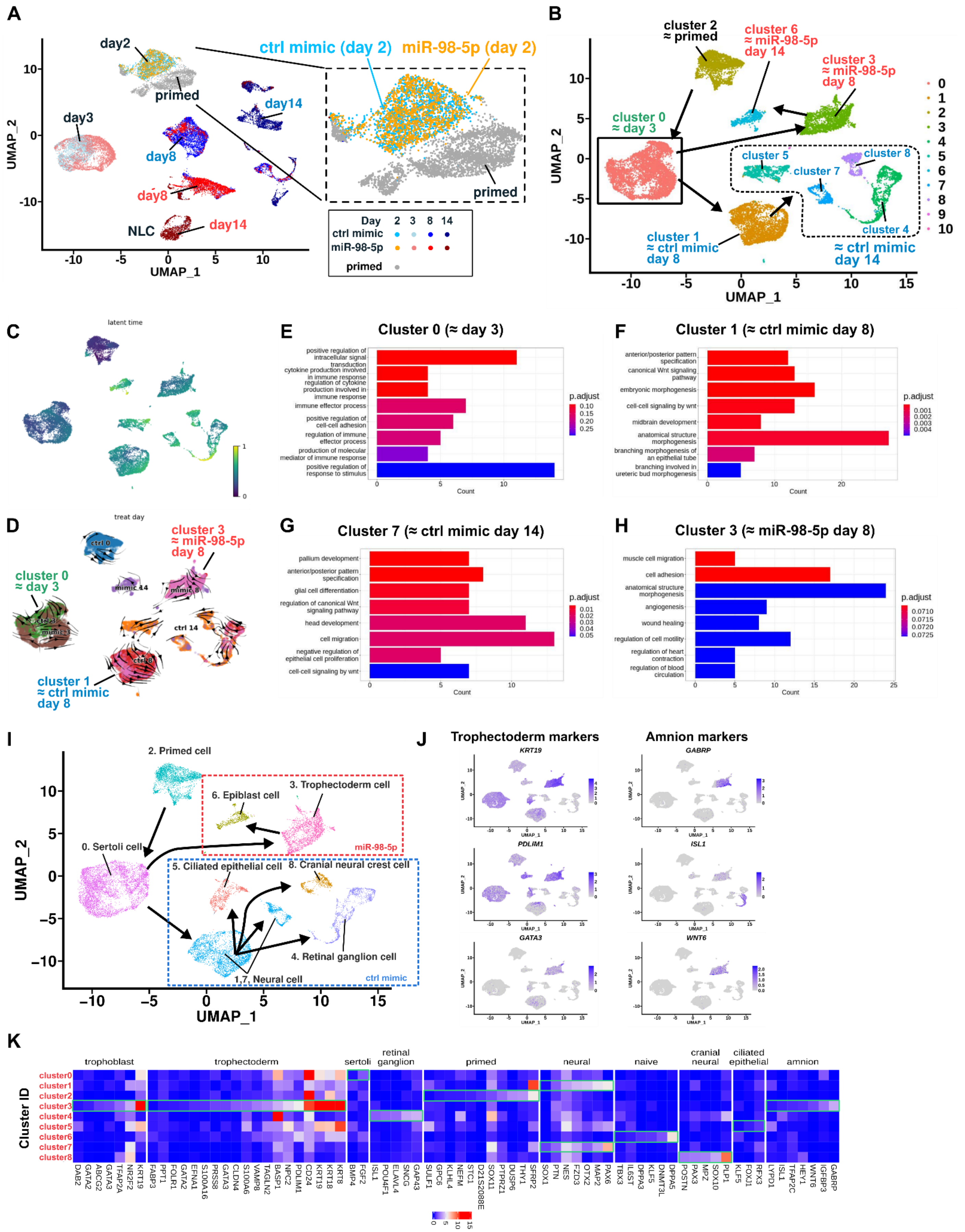


#### Supplementary Figure 3. Assignment of cell type to each cluster in scRNA-seq analysis

A: UMAP analysis of gene expression in primed cells and cells at different time points including day 2 (days 2, 3, 8, and 14) during resetting. The distribution of primed and control- or miR-98-5p mimic-transfected cells on day 2 is enlarged and displayed on the right.

B: Clustering analysis of cells (primed, days 3, 8, and 14). Almost all cells harvested on day 3 were clustered into Cluster 0. Since very few cells belonged to Clusters 9 and 10, their annotation was omitted.
C: UMAP colored by velocity-inferred latent time. Cells progress from early (purple) to late (yellow) states, consistent with actual sampling days.
D: Velocity stream plot overlaid on UMAP, colored by treatment day. Cluster 0 (≈ day 3) shows two flow directions, suggesting early divergence. Clusters 1 (≈ ctrl mimic day 8) and 3 (≈ miR-98-5p day 8) also exhibit mixed flows, indicating heterogeneous state transitions under different treatments.
E, F, G, H: Gene ontology analyses of the top 50 upregulated genes in Clusters 0, 1, 7, and 3 respectively, compared to other clusters. Any genes with a padj value higher than 1e-50 were excluded from the analysis.
I: Automatic cell type assignment by ScType[1] using marker genes obtained from the cell marker database, CellMarker 2.0[2]. Cluster numbers before the cell name are the same as cluster IDs in B. Cluster 2 is defined as primed cells because the cells in cluster 2 are H9EOS cultured and maintained according to a feeder-free protocol. Cell types in other clusters are named from cells with the highest total "cell type score" of each cell in the cluster. Clusters 9 and 10 are omitted because of few cells.
J: Global expression level of typical trophectoderm and amnion markers.
K: Mean expression of related genes of cells shown in each cluster (g0–g8 correspond to Cluster 0–Cluster 8) in A. Regions surrounded by light green squares in the heatmap show the assigned cell type gene expression for each cluster in F. Genes of trophoblast and trophectoderm are selected from references,[3,4] respectively.

#### Supplementary Figure 4

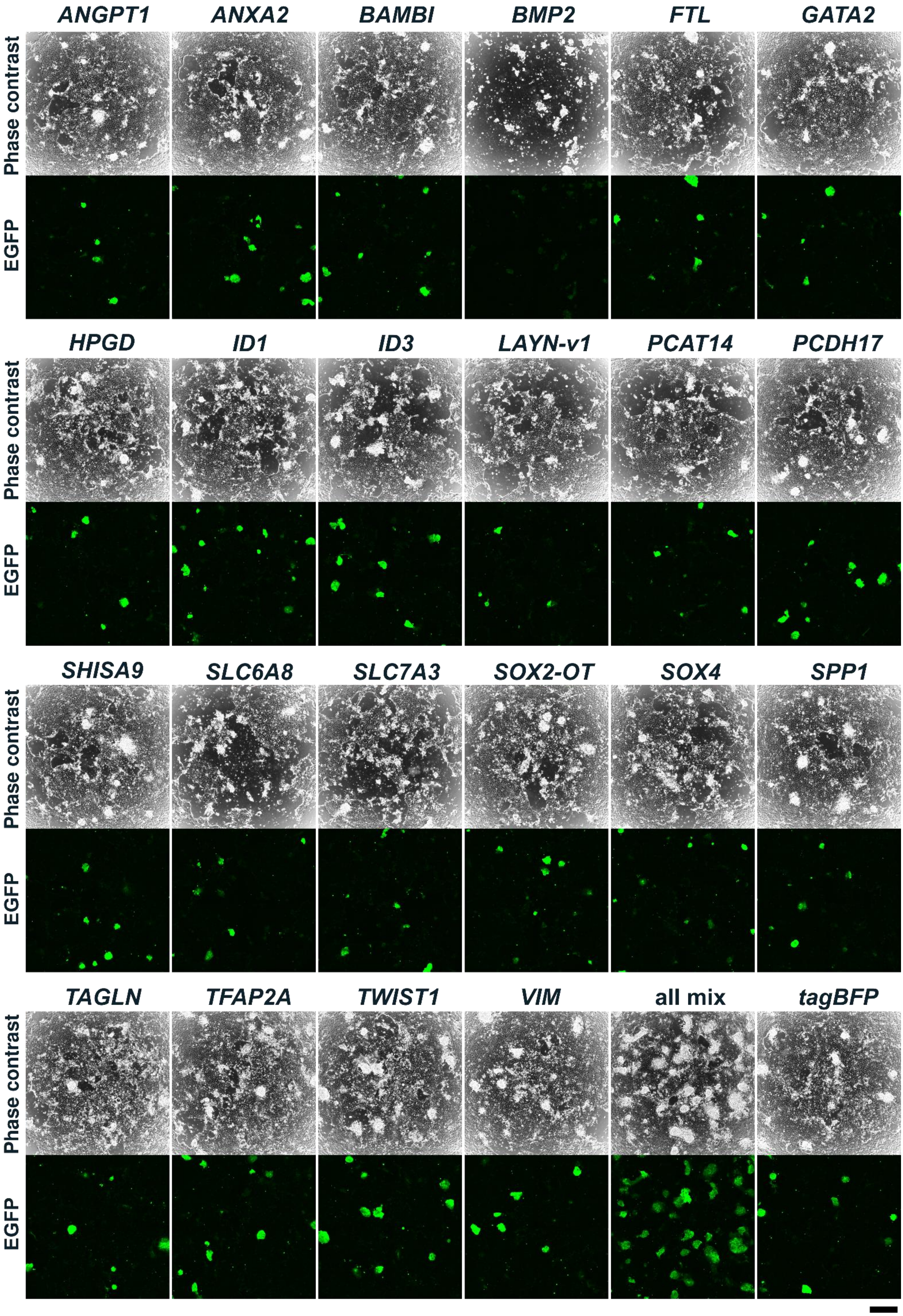

**Supplementary Figure 4. Effects of overexpression of candidate genes on resetting efficiency**

Effects of overexpression of candidate genes upregulated by miRNAs at day 3 on resetting efficiency. Candidate genes induced by doxycycline-responsive plasmid are shown as gene symbols. TagBFP was used as a transfection control. Plasmid transfection on day 1, followed by doxycycline induction for 3 days. Fluorescent images were captured on 14. Scale bar = 500 μm

#### Supplementary Figure 5

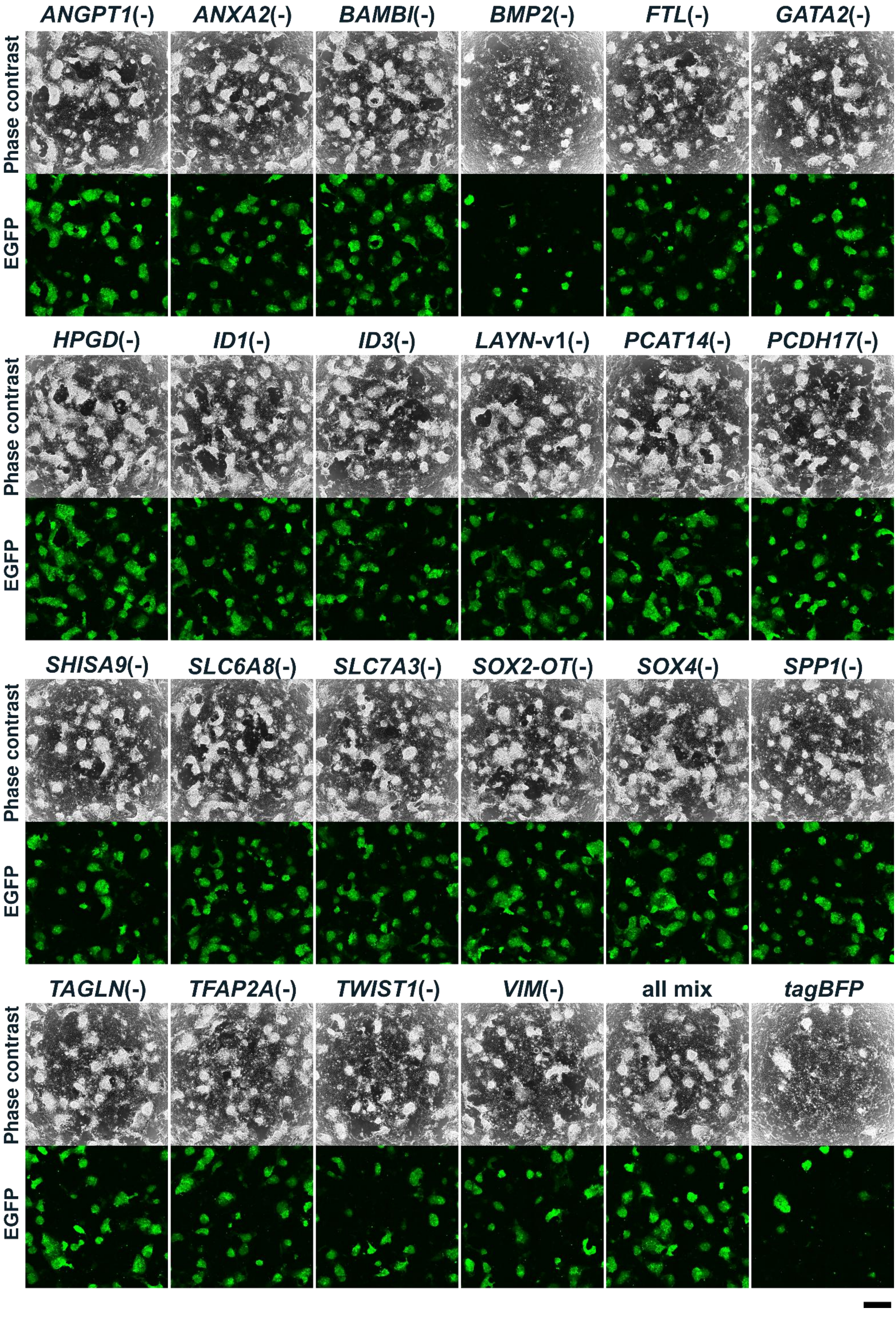

**Supplementary Figure 5. Effects of removing candidate genes from gene cocktail on the resetting efficiency**

Effects of overexpressing of mixtures of candidate genes upregulated by miRNAs at day 3 on the resetting efficiency. Cells were co-transfected with mixtures of plasmids except for 1 of the 22 candidate genes plasmids, identified by its gene symbol. TagBFP was used as a transfection control. Fluorescent images were captured on day 14. Scale bar = 500 μm.

### Supplementary Figure 6

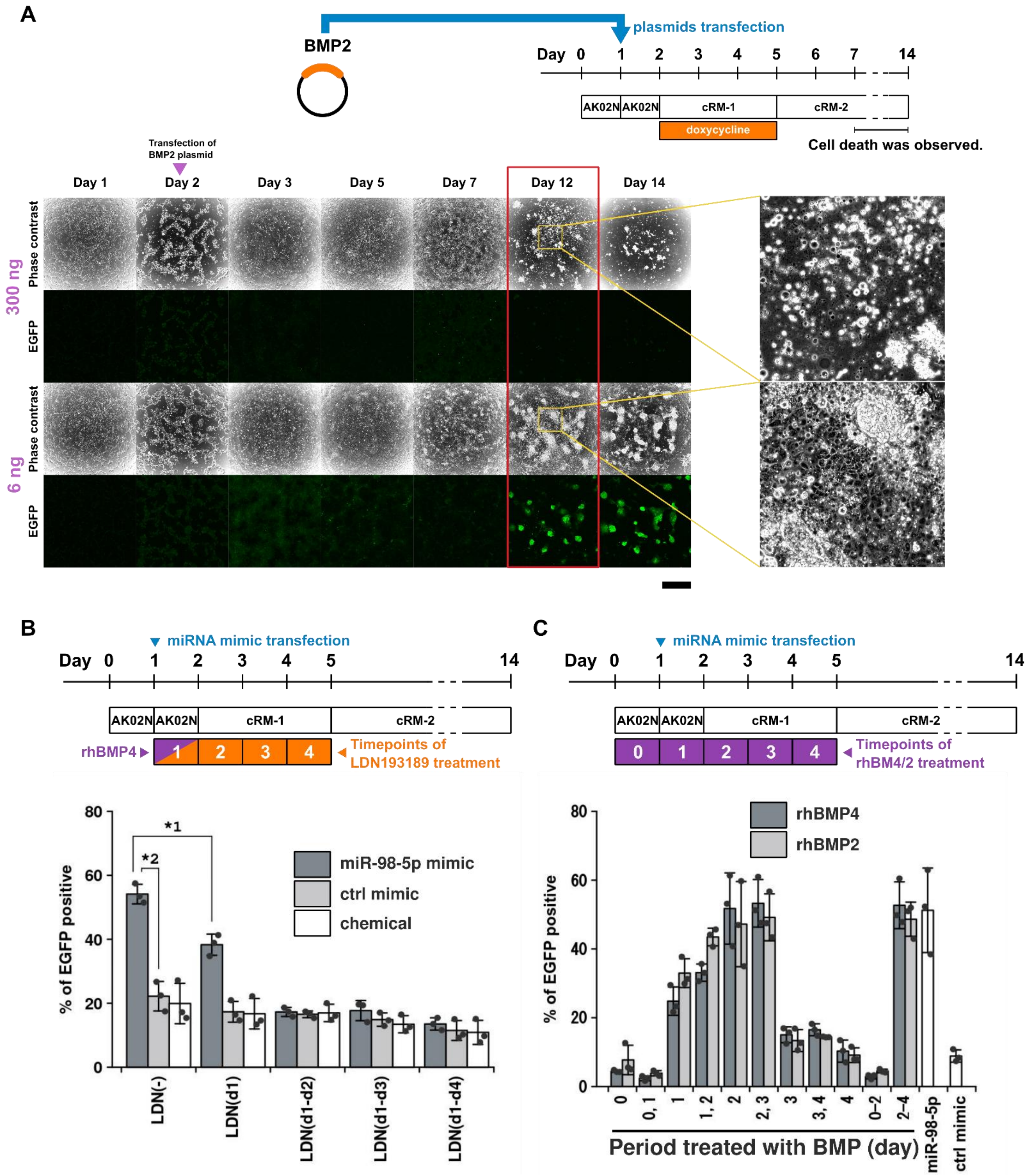


**Supplementary Figure 6. Effects of BMP signaling at various times on resetting efficiency**

A: Effects of BMP2 overexpression on cell survival. Time-lapse images of cells transfected with high (300 ng) and low (6 ng) quantities of a *BMP2* plasmid. High-magnification images of a selected region of a microscopic image are provided on the right. Cell death was only observed from day 7, long after transfection and induction by doxycycline treatment. Scale bar = 400 μm.

B: Effects of suppressing BMP signaling on resetting efficiency of cells transfected with miR-98-5p or control mimics. The percentage of EGFP-positive cells on day 14 when miR-98-5p transfected cells were cultured in the presence and absence of LDN193189. Numbers in brackets indicate the period (day) during which LDN193189 was added to the medium. *1 $P = 3.79 \times 10^{-3}$. *2 $P = 1.12 \times 10^{-3}$.

C: Comparison of resetting efficiencies in the presence of rhBMP4 and rhBMP2.The percentage of EGFP-positive cells at day 14 when cells were cultured with rhBMP4 or rhBMP2. Numbers indicate the period (day) treated with rhBMP.

#### Supplementary Figure 7

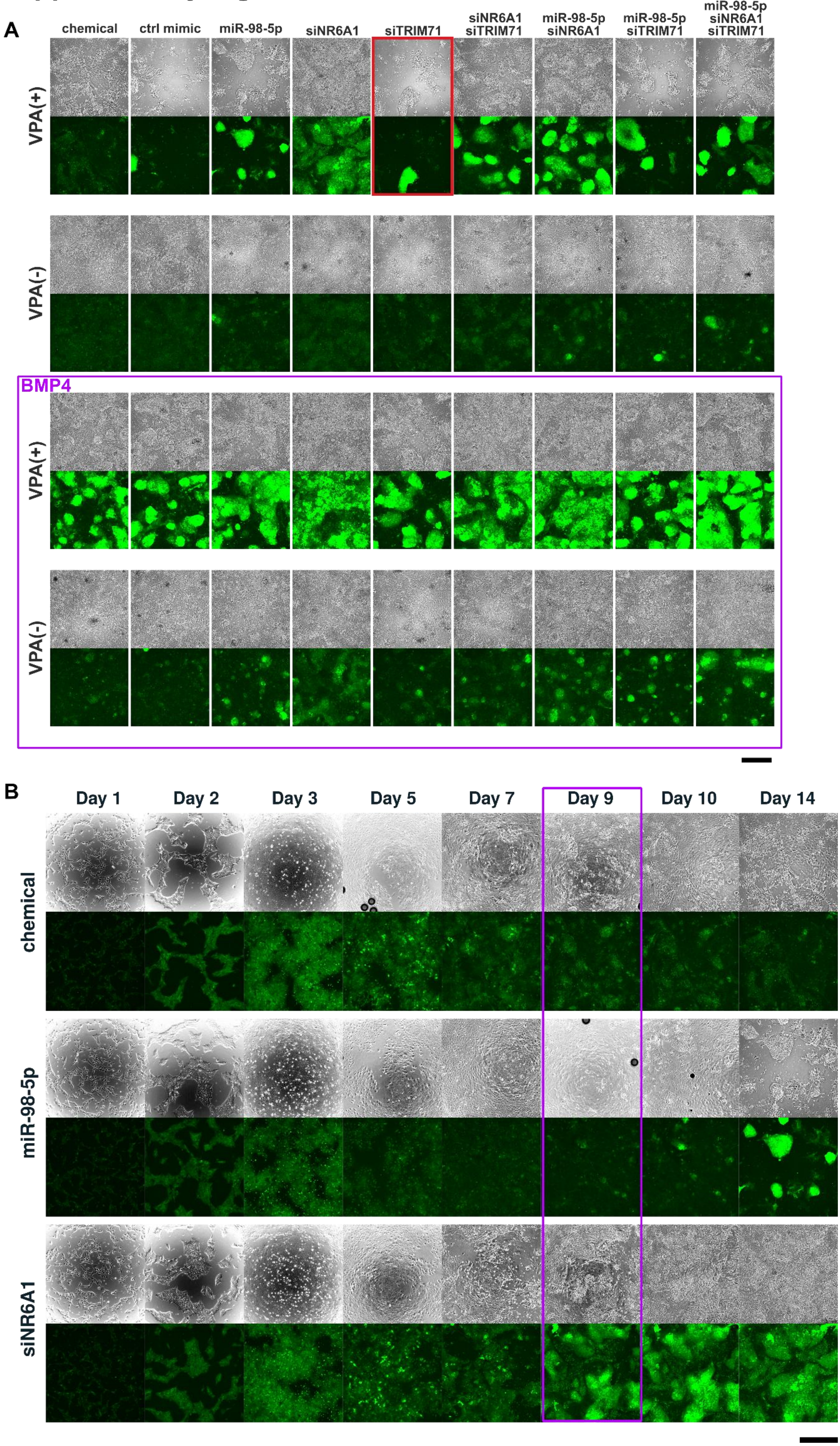

**Supplementary Figure 7. Fluorescent images on day 14 after treatment with the combination of siRNA (siNR6A1 and siTRIM71), miR-98-5p, and rhBMP4**

A: Fluorescent images on day 14 after treatment with the combination of siRNA (siNR6A1 and siTRIM71), miR-98-5p, and rhBMP4. A red rectangle indicates cells treated with siTRIM71 and VPA. Scale bar = 500 μm.

B: Time-lapse images of cells transfected with miR-98-5p and siNR6A1. Cells transfected with siNR6A1 on day 1 showed EGFP fluorescence earlier than other cells. Scale bar = 500 μm.

### Supplementary Figure 8

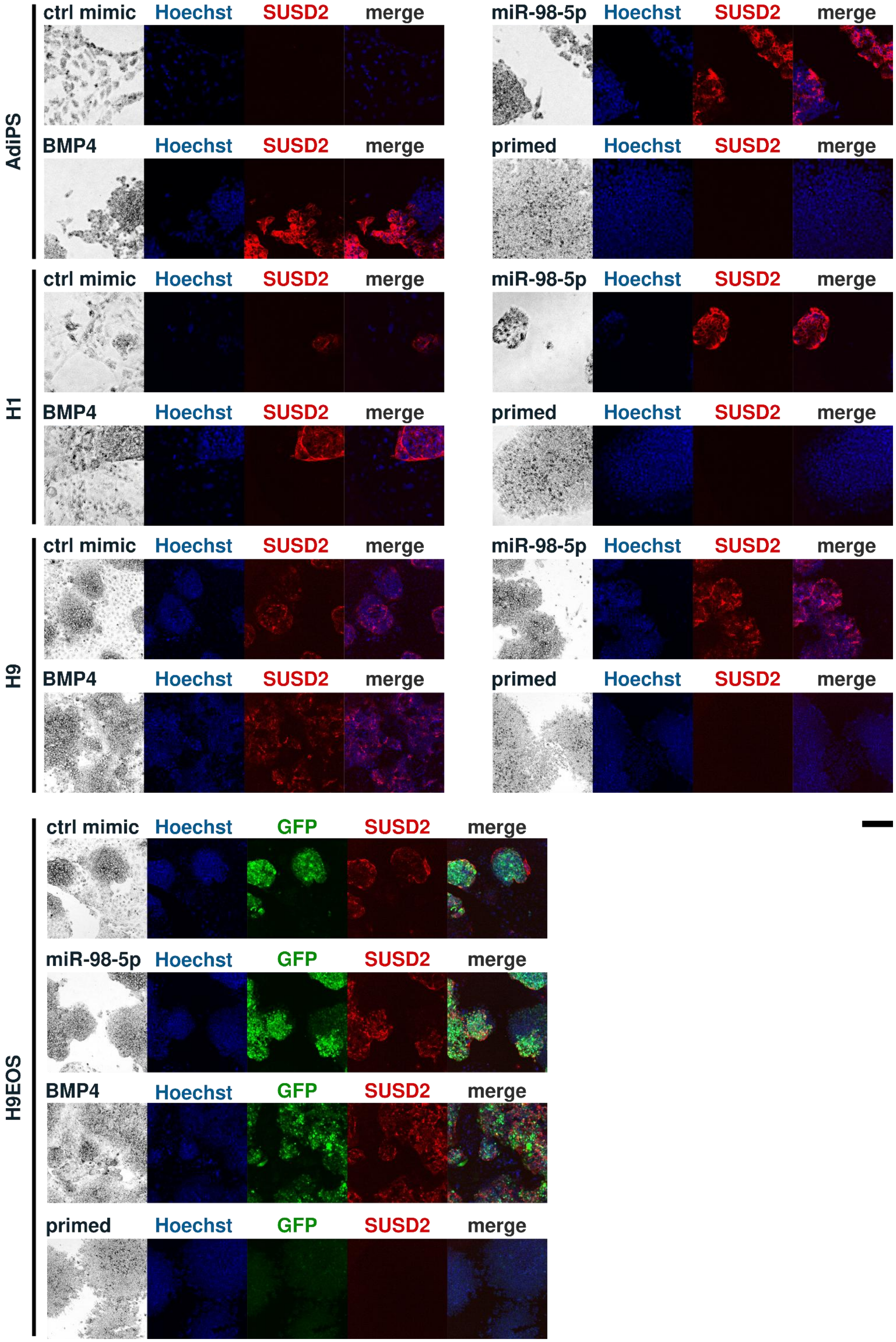

**Supplementary Figure 8. Fluorescent images of Immunofluorescent staining of SUSD2 of ESCs and iPSCs**

Immunofluorescent staining of SUSD2 in AdiPS, H1, H9, and H9EOS cells on day 14 after treatment with miR-98-5p or rhBMP4. Images are maximum-intensity projections derived from confocal fluorescent images. Scale bar = 200 µm.
